# Multiple context discrimination in adult rats: sex variability and dynamics of time-dependent generalization of an aversive memory

**DOI:** 10.1101/2024.10.06.616911

**Authors:** Fernanda Nogueira Lotz, Kétlyn Talise Knak Guerra, Ana Paula Crestani, Jorge Alberto Quillfeldt

## Abstract

Memory generalization involves the transfer of conditioned fear responses to novel contexts, a phenomenon observed in systems consolidation, whereby a time-dependent reduction in discrimination precision occurs due to the reorganization of brain regions supporting memory retrieval. To understand the fine temporal structure of this process across sexes, young adult female and male rats were trained in contextual fear conditioning and tested in the same or one of three distinct novel contexts at 2, 28 or 45 days post-training. Neutral contexts were designed to allow graded levels of fear expression relative to the training context, and sex differences were evident at the recent memory test. This pattern, however, disappeared over time due to partial generalization, with fear converging into similar, higher values, grouped into two levels for both sexes. In all experiments, females were better discriminators and displayed lower fear responses than males, apparently prioritizing different sensory modalities, with multivariate analysis suggesting that chamber size was salient for females and floor texture for males. This study is the first to compare fear responses between adult female and male rats across multiple neutral contexts and timepoints revealing several dimorphic findings.

## INTRODUCTION

The ability to store and retrieve information is a highly adaptive, evolutionarily conserved trait in animals, and in many situations the ability to predict the safety or threat of future events overcomes the capacity to retain more precise memories of past events, which is understood as an adaptive advantage (Asok et al. 2019b; Suddendorf 2006). This is especially true of fear memories in which the generalization of threat cues allows animals to increase their survival through the transference of learned behaviors to these new and potentially dangerous environments or situations based on similar cues (Bergstrom 2020; Young 2012).

Mammalian brains appear to have evolved to alternate between discrimination and generalization when encountering novel situations. However, the exact boundary conditions that determine whether a slightly distinct context triggers one outcome or the other are not completely understood. To what extent feature detection relies on a specific sensory modality, or a combination of modalities, remains unclear, and it can be difficult to isolate which elements – e.g., floor texture, wall shape, chamber size, room, size, chamber shape or cleaning agent scent – actually support graded levels of contextual discrimination, enabling measurable, statistically significant increments/decrements in responses, resembling, for instance, a “staircase” pattern (Burn 2008; Huckleberry et al. 2016; Wotjak 2019; Giachero et al. 2024).

While aversive memory generalization can be of adaptive value, overgeneralization - an exaggerated fear response to neutral cues or contexts - can have maladaptive consequences, such as restricting important resource-seeking behaviors or even inducing mental disorders as seen in humans. Therefore, to maximize the possibility of survival and successful offspring rearing, there has to be an optimal balance between threat avoidance and safety perception. This form of fear discrimination can also be understood as learned fear inhibition in the presence of safety signals whereas generalization may arise from impaired safety signaling (Day and Stevenson 2020).

In humans, excessive fear generalization is a hallmark of anxiety and trauma-related disorders such as posttraumatic stress disorder (PTSD) and generalized anxiety disorder (GAD), with women being at disproportionately higher risk of developing these conditions (Besnard and Sahay 2016; Jasnow et al. 2017; Keiser et al. 2017; Lissek and Grillon 2012; McLean et al. 2011), suggesting an involvement of sex steroids in this process. While different behavioral phenotypes are nothing new, only recently has there been an increase in the awareness of the importance of symmetrically including subjects of both sexes in preclinical and clinical studies to avoid overlooking important differences (e.g., NIH Policy on Sex as a Biological Variable, 2016).

Despite the fact that sex hormones are known to interfere with spatial, social and fear memory formation (Taxier et al. 2020), discrimination/generalization studies have produced mixed results, with some showing females generalizing before males (Lynch et al. 2013, 2014), while others indicating the exact opposite (Crestani et al. 2022; Foilb et al. 2018), or simply finding no difference between sexes (Clark et al. 2019; Keeley et al. 2015; Reppucci et al. 2013). This underscores the importance of elucidating the experimental parameters and contextual configurations that could explain contradictory findings, i.e., knowing the extent to which these might rely on sex differences and/or sensory modality, and how this allows a graded discriminatory response. In other words, knowing how specific cues present in a neutral, supposedly safe context, can trigger the aversive memory, either partially or completely, and to which extent cues signaling safety can effectively induce learned fear inhibition (Asok et al. 2019a), or whether they only undergo time-dependent generalization.

In rodents, the retrieval of an aversive contextual memory depends, initially, on the hippocampus, but, with time, becomes more reliant on neocortical structures, a process known as *Systems Consolidation* (Dudai 1996; Haubrich et al. 2016). Evidence suggests this process develops in two interrelated dimensions - a gradual neuroanatomical reorganization, and a progressive loss of the ability to discriminate between contexts, in which animals display, for instance, fear in contexts not previously paired with the aversive stimulus (Cullen et al. 2015; De Oliveira Alvares et al. 2012; Quillfeldt 2019; Rudy 2009; Wiltgen et al. 2010), a contextual memory generalization that progresses with the increasing interval between training/conditioning and test (Wiltgen and Silva 2007).

In this work, we trained young adult female and male rats in the contextual fear conditioning task and tested them 2, 28 or 45 days later, either in the conditioning context (A) or in any of three novel/neutral contexts aiming to obtain graded responses according to the different degrees of similarity in relation to the training context. To our knowledge, no previous study has compared fear responses between adult female and male rats in *multiple neutral contexts,* both in recent and remote test sessions.

## RESULTS

### PILOT STUDY

Subjects of both sexes were fear conditioned to context A, and, 2 days later, retrieval tests were performed in the same or in an alternative set of neutral contexts (P1, P2, P3 and P1’ **Figure 1.A**), sorted according to distinct intuitive criteria – at first, mostly defined by chamber shape and floor texture (see Supplemental material for a more detailed discussion on this point).

**Figure 1.**
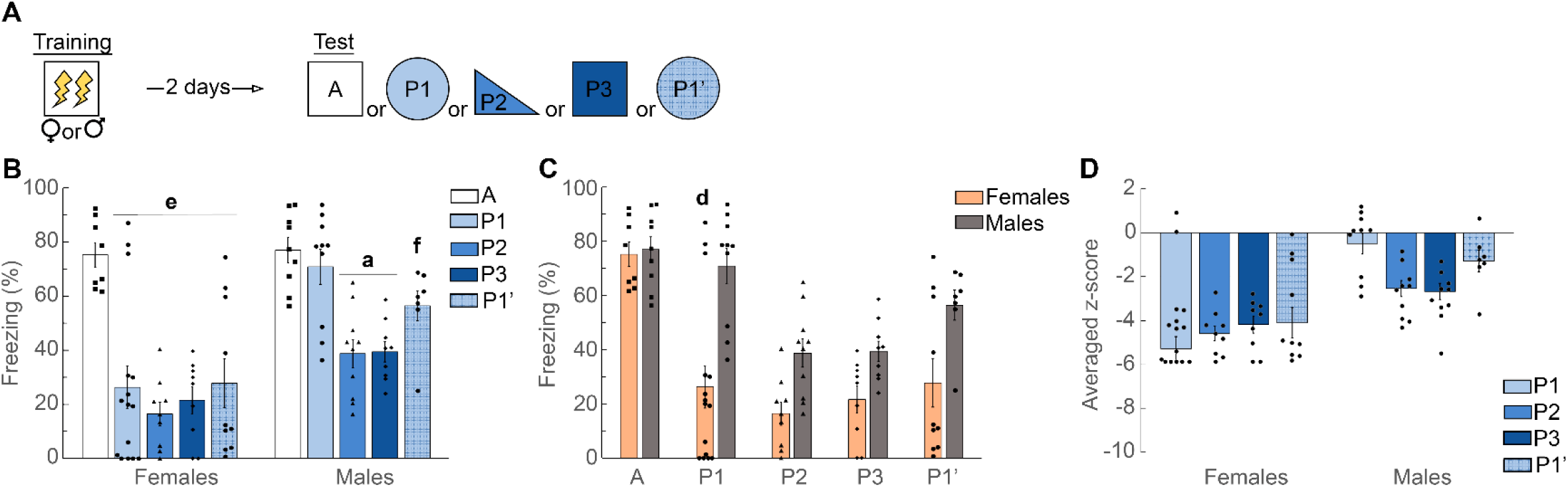
Contextual discrimination of female and male rats trained in a fear conditioning task and tested 2 days later. (**A**) <u>Experimental protocol</u>: all subjects were trained in Context A and subsequently tested either in A, P1, P2, P3 or P1’ 2 days after training (P1’ being a slight modification of context P1); (**B**) <u>Histogram</u> showing mean±SEM of the % freezing time in each context grouped by sex, with individual dots overlayed representing individual animals; (**C**) Same data (and statistical analysis) grouped by context. B & C: Data analyzed by independent two-way ANOVA followed by Tukey *post hoc* test: **a** (different from A / P1), P < 0.0055, **f** (different from P2 / P3), P < 0.032, **d** (females differ from males tested in the same context), P < 0.0001, and **e** (all groups differ from A), P < 0.0001; (**D**) Individual (and the average) <u>Z-scores</u> for contexts P1, P2, P3 and P1’ (standard deviations, all compared to A).

Independent two-way ANOVA of female and male rats tested in one of a new set of four contexts (see **Figure S1.A**) plus a version of P1 without wood shavings on the floor (Context P1’) revealed significant effects of Context x Sex Interaction (F_(4, 86)_ = 3.059; P = 0.0208, ω^2^ = 0.08), Context (F_(4, 86)_ = 16.44; P < 0.0001; ω^2^ = 0.39) and Sex (F_(1, 86)_ = 30.62; P < 0.0001, ω^2^ = 0.24) – **Figure 1.B**: According to the ω^2^, all detected effects were large and robust. Number of subjects per context (A, P1, P2, P3 and P1’) respectively: females, 8, 15, 9, 9, 10 and males, 9, 10, 10 and 7.

In females (**Figure 1.B**), Tukey *post hoc* test revealed significant differences between groups tested in the conditioning context A and all novel contexts (P’s < 0.0001) with large effect sizes (respectively, A x P1, g = 1.896; A x P2, g = 4.55; A x P3, g = 3.848; A x P1’, g = 2.069), though freezing levels did not differ significantly between the novel contexts P1, P2, P3 and P1’ (P’s > 0.90 - see **Table S-1**). For males (also **Figure 1.B**), Tukey *post hoc* test showed groups differed significantly between Context A and novel contexts P2 (P < 0.0001, g= 2.49) and P3 (P < 0.0001, g = 2.962), but not between A and P1 (P = 0.98), A and P1’ (P = 0.57) or P1 and P1’ (P =0.90). Also, context P1 differed from P2 (P = 0.0011, g = 1.727) and P3 (P = 0.0021, g = 1.872). Male and female groups did not differ in freezing levels in the conditioning context A (P >0.99), context P2 (P = 0.324), P3 (P = 0.675) and P1’ (P = 0.12), but females displayed less freezing than males in context P1 (P < 0.0001, g = 1.656) (**Figure 1.C**).

In order to estimate the degree of discrimination, we plotted the individual and average Z-scores for every neutral context (P1, P2, P3 or P1’). For females, all data points have significantly lower values compared to A, showing that females were better discriminators across all the novel contexts compared to males (**Figure 1.D**). Also different from males, females were able to discriminate between the conditioning context and P1 (and its modified version P1’), while males displayed high levels of fear in these contexts.

### MAIN EXPERIMENT

Subjects of both sexes were fear conditioned to context A, and 2, 28 or 45 days later, different groups were submitted to retrieval tests in this or three other neutral contexts (B, C and D, **Figure 2.A**), also sorted according to distinct intuitive criteria (see Discussion and Supplemental material for a more detailed discussion).

**Figure 2.**
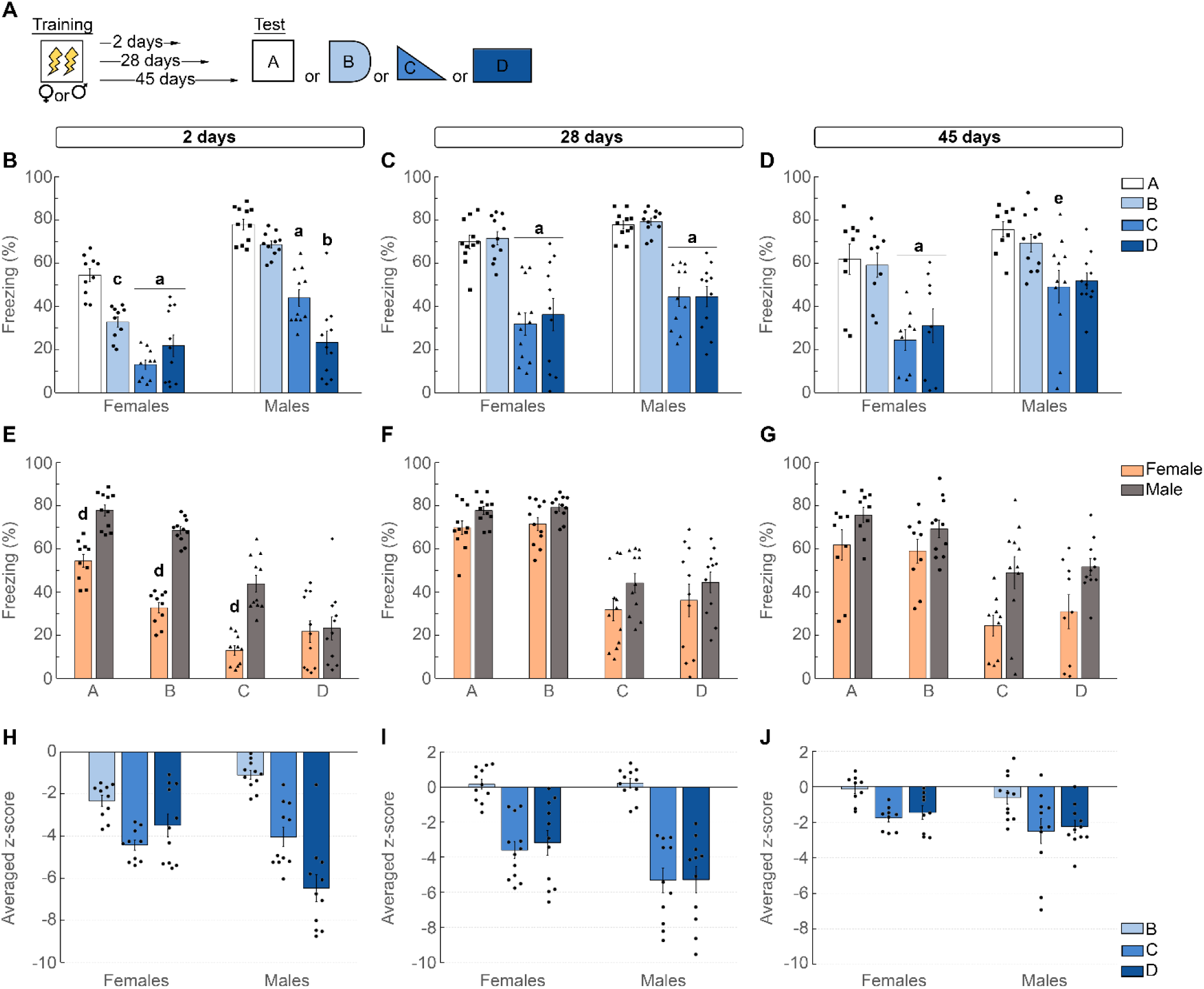
Context discrimination of female and male rats 2, 28 or 45 days after training in a fear conditioning task. (**A**) <u>Experimental protocol</u>: all subjects were trained in Context A and subsequently tested either in A, B, C or D 2, 28 or 45 days later; (**B, C and D**) <u>Histograms</u> showing mean±SEM of the % freezing time in each context grouped by sex with individual dots overlayed representing individual animals; (**E, F and G**) Same data (and statistical analysis) grouped by context. B - G: data analyzed by independent two-way ANOVA followed by Tukey *post hoc* test. **a** (different from A / B), P < 0.015, **b** (differs from same sex groups tested in all other contexts), P < 0.002, **c** (different from A), P = 0.002, **d** (females differ from males in this specific context E), P < 0.0005, and **e** (different from A), P = 0.03; (**H, J and I**) Individual (and average) <u>Z-scores</u> for contexts B, C and D (standard deviations compared to A).

When animals were tested 2 days after training in one of four distinct contexts (See **Figure 4.B**), independent two-way ANOVA revealed significant effects in the interaction between Context x Sex (F_(3, 77)_ = 9.182; P < 0.0001, with effect size ω^2^ = 0.22), Context (F_(3, 77)_ = 63.92; P < 0.0001, ω^2^ = 0.699) and Sex (F_(1, 77)_ = 82.17; P < 0.0001, ω^2^ = 0.49) (**Figure 2.B, 2.E and 2.H**): according to the ω^2^, all detected effects were large and robust. Number of subjects per context (A, B, C and D respectively): females, 10, 10, 10, 11, males, 11, 11, 11, 11. In females, Tukey Multiple Comparisons *post hoc* test revealed different freezing levels in contexts A and B (P = 0.002, g = 2.256), but not between contexts C and D (P = 0.67); in males, there are significant differences between A and C (P < 0.0001, g = 3.132) and A and D (P < 0.0001, g = 3.923), B and C (P = 0.0001, g = 2.483) and C and D (P = 0.002, g = 1.331), but not between A and B (P = 0.572) (**Figure 2.B** and see **Table S-1**).

For rats tested at 28 days in one of four distinct contexts, independent two-way ANOVA revealed no significant effect in the interaction between Context x Sex (F_(3,81)_ = 0.132; P=0.94), while Context (F_(3,81)_ = 43.81; P < 0.0001, ω^2^ = 0.62) and Sex (F_(1,81)_ = 8.65; P < 0.0043, ω^2^ = 0.09) differ significantly (**Figure 2.C, 2.F and 2.I**): according to the ω^2^, effect size was just moderate to large for Sex, while for Context they were large and robust. Number of subjects per context (A, B, C and D respectively): females, 11, 11, 12, 11, and males, 11, 11, 11, 11. For both females and males, Tukey Multiple Comparisons *post hoc* test revealed similar freezing levels in all contexts, with no differences observed between contexts A and B (P > 0.99) and between C and D (P > 0.99) (**Figure 2.C** and see **Table S-1**).

Finally, for animals tested 45 days after training, independent two-way ANOVA revealed no significant effect in the interaction between Context x Sex (F_(3,70)_ = 0.6543; P = 0.583), while Context (F_(3,70)_ = 15.52; P < 0.0001, ω^2^ = 0.36) and of Sex (F_(1,70)_ = 18.27; P < 0.0001, ω^2^ = 0.18) differ significantly (**Figure 2.D, 2.G and 2.J**): according to the ω^2^, all detected effects were large and robust. Number of subjects per context (A, B, C and D respectively): females, 9, 9, 9, 9, and males, 9, 11, 11, 11. Tukey Multiple Comparisons *post hoc* test revealed that females maintained a profile similar to that of 28 days, with no significant differences between A and B (P > 0.99) or C and D (P = 0.99), while males keep discriminating between contexts A and C (P = 0.029, g = 1.348), but not between A and D (P = 0.07). Also, when compared in the same context, there were no differences between males and females (P > 0.05 for all comparisons - see **Table S-1**) (**Figure 2.D**).

In order to estimate the degree of discrimination, we plotted the individual and average Z-scores for every neutral context (B, C or D) compared to the conditioning context A. For rats tested at 2 days (**Figure 2.H**), all data points have lower values compared to A, and they differ among them, except for context C, that seems to be equally well discriminated by both sexes. Importantly, the salient “staircase effect” in the degree of discrimination of the males was a sorting pattern not followed by the females in these contexts. For animals tested at 28 days (**Figure 2.I**), the main effect was the dramatic response to context B – now indistinguishable from A – for both females and males, while contexts C and D converged to a similar degree of discrimination: in sum, females appear to better discriminate these contexts than males. The same convergence of responses (% freezing level) between contexts B and A observed at 28 days persisted at 45 days (**Figure 2.J**). Discrimination of contexts C and D in males, however, was much less evident now, suggesting that, at this timepoint, all neutral contexts tended to evoke a fear response similar to A, with average z-scores falling within −0.6 and −2.5 standard deviations for males, and −0.131 and −1.760 for females.

## DISCUSSION

We initially selected multiple novel contexts based on intuitive criteria, with the goal of obtaining graded levels of fear expression, i.e., each one a statistically significant reduction in the freezing levels in relation to the conditioning context. Then, we observed and quantified eventual changes in freezings over time to investigate the fine temporal structure of systems consolidation in rats of both sexes. This phenomenon is characterized by a time-dependent fear increase in novel contexts, usually interpreted as memory generalization. With this experimental setup we aimed to verify how generalization to distinct contexts evolves over time in female and males, either continuous or discontinuously.

There is no clear definition regarding which components of a behavioral context are preferred by rodents, and even less is known about preferences in relation to sex (Burn 2008; Huckleberry et al. 2016; Wotjak 2019; Giachero et al. 2024). In planning our *graduated contexts*, instead of simply varying one or two sensory criteria, as usually done (Shepard 1987; Young 2012), we tried an intuitive, naturalistic *approach* that respected the multidimensional nature of the task, since discrimination between contexts involves the integration of several sensory modalities simultaneously. **Figure S-1** summarizes several of these parameters in the selected contexts, and while clearly not an exhaustive list, it aims to capture the more salient sensory modalities - such as olfactory, tactile or visual features.

### SORTING OF MULTIPLE CONTEXTS: RECENT MEMORY VS. SEX

As mentioned above, we attempted to obtain graded responses with a first set of neutral contexts - P1, P2, P3 and P1’ – but were not very successful (**Figure 1.B**). There was a noticeable difference between sexes, with females discriminating context P1 from A, while males did not. This was surprising because P1 looked, intuitively, to be the least similar context to the conditioning one. Since context P1 had wood chips spread on its smooth floor, we suspected this could have acted as a reminder of their homecage (in which wood chips were used as bedding), somehow signaling safety in an otherwise very different environment - or even acting as a distractor. Therefore, we tested the animals in a cleaner variant, P1’, devoid of wood chips, but the freezing response of males remained higher to this context, while females had, again, lower levels, discriminating it from A just as they had with P1. In sum, females appear to be better discriminators than males at least for recent memory, distinguishing each of the four neutral contexts used in the pilot study equally well, all exhibiting a significantly lower freezing compared to the fear response in the conditioning context A. This dimorphic behavioral response suggests each sex prioritizes different parameters in distinct ways.

It is possible that the abnormally higher fear response of males in such dissimilar contexts P1/P1’ just two days after training might be due to a novelty-induced effect (Maren 2014; Greggor et al. 2015). We did not pursue this avenue of investigation, but in order to avoid this kind of interference in recent memory and verify which contextual features might endure across a time-dependent generalization, a different set of neutral contexts was chosen for the main experiment (B, C and D – see **Figure S-1**) in order to compare female and male fear responses at different training-test intervals.

These contexts, despite also intuitively sorted based mainly on chamber size and cleaning agent scent, allowed us to obtain a clear series of decreasing freezing values in males (see **Figure 2.B**, right panel). Females, however, displayed a completely different pattern of fear expression, evincing again a difference between sexes (**Figure 2.B**, left panel). We intentionally pursued a "staircase-like" aspect observed in male freezings in contexts A, B, C and D, but females also displayed a graduated set of responses, only in a different order – A, B, D and C – and with a distinct statistical robustness. Females and males definitely exhibit distinct behavioral responses when exposed to multiple contexts.

Employing the above protocol, with this selection of contexts, high levels of freezing were obtained in the conditioning context A in all training-test intervals, indicating *good acquisition (learning) and retention* of the aversive experience (**Figures 2.B, C** and **D**).

Of note, two days after training, males were definitely *not able to distinguish* context B from A, while females displayed a statistically significant lower freezing level in B compared to A (c in **Figure 2.B** and d in **2.E**). This sex asymmetry is no longer observed at 28 days, when B becomes indistinguishable from A for both females and males (**Figures 2.C** and **2.F**), persevering up to 45 days (**Figure 2.D** and **2.G**). A study by Huckleberry and colleagues (2016) using male mice tested how different sensory modalities - scent, floor texture or chamber roof shape, or a combination of all three, influenced the generalization of fear to a novel, modified version of the conditioning context. They found that floor texture and scent were more salient than predominantly visual cues (roof) to produce a higher fear response in the novel context. Despite having some control upon context design, this work only covers recent memory (1-2 days), but the main problem is that it was done in mice (and males only), and several behavioral differences are known to exist between these species.

In our recent memory test, we introduced context B which is a modified version of the original conditioning chamber A, with the grid covered by a smooth linoleum floor, a curved wall added inside the chamber and cleaned with the same solution as in A (Ethanol). Nevertheless, based on how males reacted, we cannot discard the possibility that, for them, the same odor from the original training context A was inadvertently detected in B (reinforced by the fact that the same cleaning agent – ethanol - was used in A and B), notwithstanding the other modified parameters (inner chamber shape and floor texture). The higher fear response of male rats could be due to the *transference* of enough aversive information from A to B. However, if this is true, considering that females were able to adequately discriminate between these contexts, suggest that males, somehow, value olfactory cues more than females, at least when testing recent memory. This is consistent with several studies that show the prominence of olfactory threat signals in rodent life (Canteras et al. 2008; Grella et al. 2020; Takahashi et al. 2005).

Another noticeable fact is that, in the main experiment recent memory test, females express *a lower maximum freezing level* compared to males not only in context A, but also in B and C (**Figure 2.B**), but not in D - the least similar, in our view (**Figure 2.E** and **Figure S-1**), pointing again to a superior discriminative capacity of young adult females relative to males, a general difference in fear expression that could be considered another sexual dimorphism if not for the fact that in the Pilot experiment this difference was not observed (**Figure 1.B**), possibly due to seasonal variability of rat behavior over a span of months. Nonetheless, lower fear in females compared to males is consistent with some studies comparing conditioned fear between males and females (Colon et al. 2018; Morena et al. 2021; Trott et al. 2022b; Soares et al. 2024; Vazquez et al. 2024), but might differ from other findings using a distinct aversive task (Lynch et al. 2013).

Figure 3, below, schematically summarizes most of the similarities and differences of fear responses between the Pilot study and the Main experiment in multiple contexts 2 days after training.

**Figure 3.**
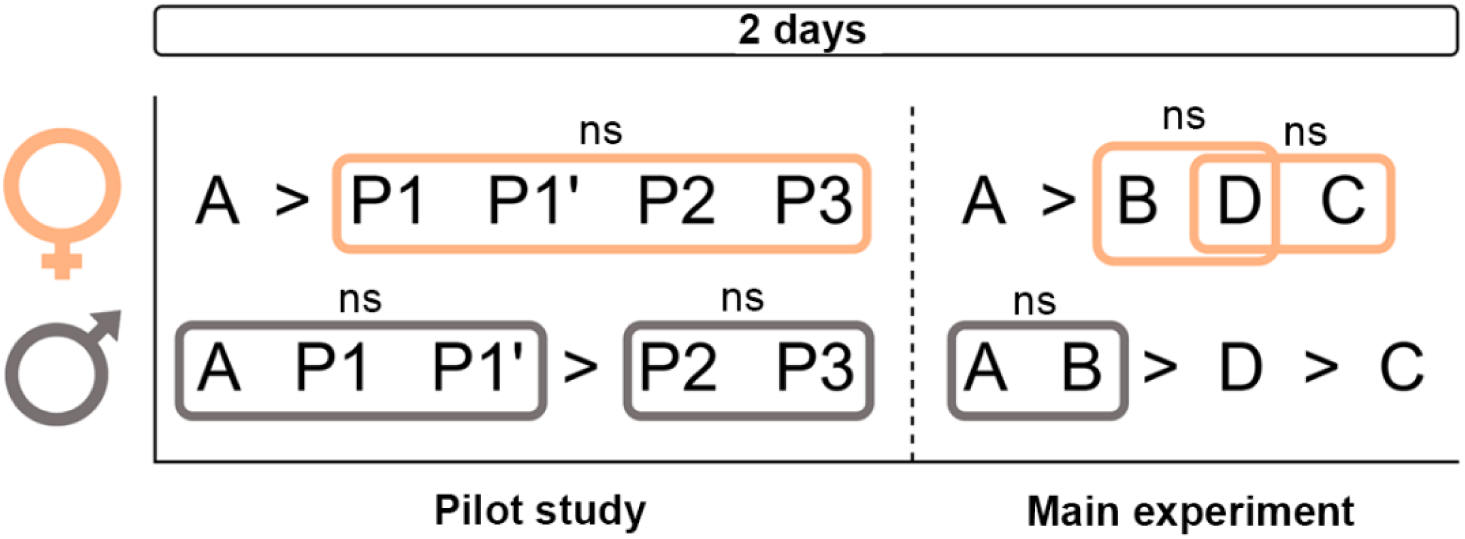
Graphic summary of the behavioral response of females and males to each context in both experiments (recent memory only). All subjects were equally trained in Context A (3 min habituation plus two 0.5 mA / 2 sec footshocks within 30 sec intervals) and, 2 days later, tested either in A, P1, P2, P3 or P1’, or in A, B, C or D (Main Experiment). “>“ or “<”, statistically significant difference, “ns”, non-significant difference.

### TIME-DEPENDENT PARTIAL OR COMPLETE GENERALIZATION OF REMOTE FEAR MEMORIES

In murine fear conditioning models, the most frequent indicator of poor context discrimination is the expression of freezing in neutral contexts in levels close to that of the conditioning one. This aligns well with the definition of generalization as the “transference of conditioned responding to stimuli that perceptually differ from the original one” (Bergstrom 2020), but does not provide enough information on *how much* “conditioned responding” is actually necessary. Over time, our results show at least two degrees of generalized response, one nearly complete, and the other, partial (see **Figures 2.C** and **2.D**): while contexts A and B become statistically indistinguishable from 28 days onwards – a case of *full* or *complete* generalization – contexts C and D show statistically significant differences from the conditioning context A (and B) at 28 for both sexes and at 45 days for females (males do not distinguish D from A at 45 days), and from their previous values, at 2 days, which implies *generalization* to an intermediate degree. However, what could account for the *partial* increase in fear response? Poorer context recognition? True lower fear? Forgetting? A trade-off strategy between pursuing rewards and avoiding harm? There might be different explanations, some even mutually exclusive, but discussions on subtypes of generalization are rarely found in the literature (but see Shepard 1987; Pearce and Bouton 2001; Kehoe 2008; Young 2012). Therefore, to ensure the phenomenon is accurately represented, why not refer to it as what it appears to be, a *partial* generalization (or perhaps *undergeneralization*, or even *subgeneralization*)? Thus, assaying multiple, graded neutral contexts may be a useful approach to quantify distinct levels of generalization.

The main experiment consisted of investigating how discrimination of each context A, B, C and D (**Figure S-2**) evolves over time for both sexes, testing two instances of remote memory compared to a recent memory performance. What we observed was that 28 and 45 days after training, animals of both sexes expressed similar freezing values, grouped in two levels. At 28 days, discrimination of contexts C and D converged to a similar level – of around 50% fear expression – significantly lower (statistically) than the response to A and B (**Figure 2.C**). The grouping of contexts C and D vs. A and B effectively disrupted the multilevel pattern observed in the recent memory test, and, in particular, dismantled the “staircase-like” effect more explicitly exhibited by males at 2 days (ABCD), but also by females in another order (ABDC - **Figure 2.B**). This new pattern persisted until 45 days, with a slight performance increase in context D in males (**Figure 2.D** and **G**). Importantly, all novel contexts displayed a statistically significant increase in freezing levels at the remote tests compared to the test at 2 days, which demonstrates the partial time-dependent contextual generalization process described above.

The persistence of the partial generalization in contexts C and D from 28 to 45 days, suggests a discontinuous progression, but to shed more light on this it would be necessary to examine much older remote memories than those explored in the present experiments in order to verify if C and D converge asymptotically to the same level of freezing in A and B (i.e., a full generalization). But considering that forgetting would also be expected to increase and counterweigh the variable being measured (% freezing), this seems less probable. For now, for females at least, with the contexts we used, there was no indication that, over time, fear generalization exhibited any type of continuous increment, much less increasing until reaching the conditioning context performance level. For males, we observed statistically equivalent levels between freezing in contexts A and D at 45 days, but the same cannot be said for context C.

### MULTIPLE INSTANCES OF DIMORPHISM

Our experiments, employing two sets of multiple contexts allowed the detection of a wide variety of behavioral responses, evincing some instances of sexual dimorphism (see **Table 1**). Young adult rats exhibited a diversity of sex- and time-dependent fear responses in contexts sorted according to multisensory criteria (tactile, visual, olfactory, or their combinations), displaying at least two kinds of time-dependent transitions from precise to generalized fear memory. The diversity of responses, coupled with the challenge of accurately defining truly graded contexts may help explain the pervasive difficulties encountered by researchers in identifying suitable neutral contexts when planning experimental setups to investigate memory generalization (Bouton et al. 2021; Giachero et al. 2024; Rosas and Alonso 1997; De Oliveira Alvares et al. 2012). Any specific animal preference, including sexual differences, should be added to a much-needed list of *Systems Consolidation Boundary Conditions* (SCBC).

**Table 1**, below summarizes these dimorphic findings, most of which may (or may not) share a common cause, the investigation of which was beyond the scope of this study.

**Table 1.**
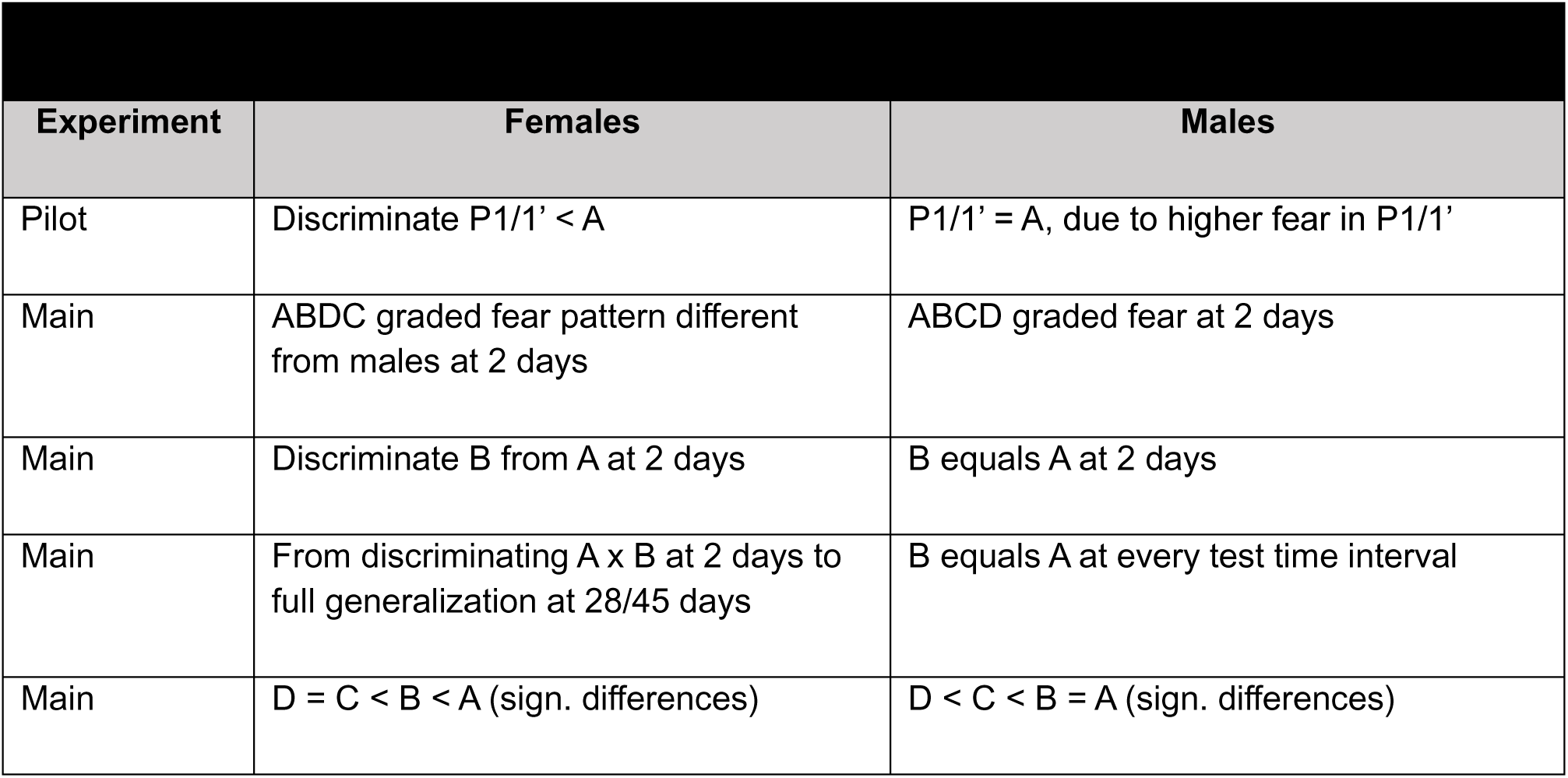
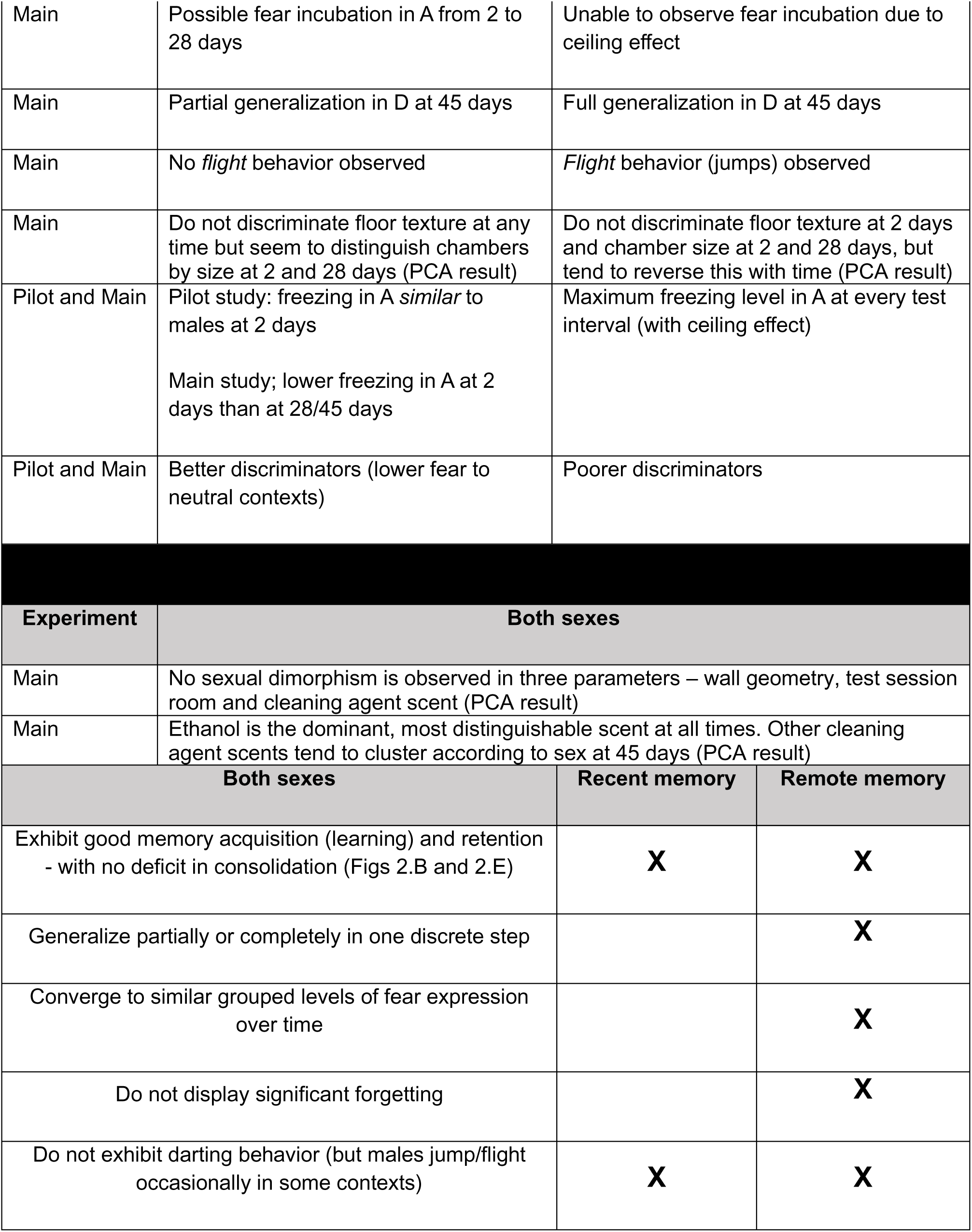
Summary of dimorphic and non-dimorphic behaviors observed in the Pilot Study and Main Experiment.

### MEMORY GENERALIZATION OR SOMETHING ELSE?

As Shepard (1987) explains, “generalization is thus a cognitive act, not merely a failure of sensory discrimination” that amounts to “inferring if an object or context belongs or not to what may be considered ‘a consequential region’, i.e., a safety “region in an individual’s psychological space”. The kind of fear generalization we try to evince here is specifically the time-dependent loss of trace precision characteristic of Systems Consolidation, i.e., the transference, with time, of a conditioned memory to novel contexts (Bergstrom 2020; Young 2012), but since our quantifiable variable is the fear response itself, we must consider that different processes could explain the observed change. Our data allow to discard at least two confounding factors, *forgetting* of the aversive experience and *fear incubation*.

It is well established that high emotional arousal such as that elicited in fear-driven tasks is important to consolidate long-lasting memories, while less emotionally charged memories tend to decay more easily with time. Here however, freezing performance in the conditioning context remained high for both sexes – over 60% of the session duration – up to 45 days post-training (see **Figures 2.D** and **2.G**), confirming that the amount of forgetting over time, if any, can be considered negligible.

In novel contexts, however, the specific contribution of forgetting to the retrieval of fear memories can be difficult to distinguish from a true generalization process. For instance, memory for contextual cues signaling safety might undergo gradual decay with time and give way to other threat signals that become more salient, e.g., the tactile perception of a metal grid floor similar to the one in which footshocks were delivered. One theory proposes that the main cause of generalization is *forgetting* of the stimulus attributes (Riccio et al. 1992), but there might be other explanations, such as *associative novelty-induced fear* (Kumaran and Maguire 2006; Maren 2014), *neophobia* (Greggor et al. 2015), or even – among the factors raised to explain avoidance generalization in PTSD animal models - *non-associative novelty-induced fear,* a kind of sensitization (Pamplona et al. 2011). With the exception of forgetting, none of these additional possibilities were directly examined here. However, as males in the Pilot study expressed high levels of fear to P1 (and its variant, P1’), deemed the least similar to A, the possibility of this resulting from a novelty-induced effect motivated us to substitute the contexts of the Pilot study for another set.

Another important confounding factor would be *fear incubation*, in which fear responses incubate, i.e., increase over time (Pickens et al. 2009; Poulos et al. 2016). While the involvement of this effect in novel contexts remains an open question, some studies suggest that high levels of fear may be due, at least in part, to this phenomenon in association with retrieval error and the flattening of the generalization curve (Poulos et al. 2016; Rosas and Alonso 1997).

The dividing line between Fear Generalization (FG) and Fear Incubation (FI) remains unclear, and experimental evidence that effectively distinguish the two is scarce in the literature. We submit an operational procedure to solve this problem in the case of contextual fear conditioning: *when a time-dependent fear increase takes place only in the conditioning context,* it could be considered FI, while if the increase occurs only *in the neutral context(s)*, it could be considered a *bona fide* case(s) of FG (Poulos et al. 2016; Wiltgen and Silva 2007). Of course, to ascertain this, the %freezing level in the conditioning context must *not be at a ceiling* level (see Quillfeldt 2018), otherwise we would not be able to observe FI. In our study, males displayed much higher levels of freezing than females in A in all training-test intervals – which prevented us from confirming FI (**Figure S-2.B**), while females, tested in the conditioning context A, showed a substantial increase in % freezing from day 2 to day 28, which then remained stable up to 45 days (**Figure S-2.A**). This suggests that, differently from males, females might exhibit some degree of fear incubation which points to another instance of dimorphism.

Notably, in females, the increase of fear in B was even steeper than in A, suggesting the interesting possibility that incubated fear might itself be generalized to neutral contexts depending on the degree of similarity to the conditioning one: generalization could be, to some extent, the “transference” of FI from the conditioning to the neutral context(s), resulting in a response indistinguishable from pure generalization as it becomes diluted by the overall increase of fear expression. This level of synergism would demand a distinct experimental setup in order to be more clearly dissected, which is not possible with our data. This was, however, outside the scope of the present study. One last point to consider is that our experimental protocol is quite mild in terms of aversiveness – at least compared to the most reliable known protocols of FI induction, which usually involve extensive training combined with multiple, high-intensity CS (Pickens et al. 2009), so perhaps FI is not to be expected.

### STRESS, FEAR AND THE LIMITS OF FREEZING RESPONSE AS A MEASURE

One additional factor that may play into sex differences in fear memory generalization is stress, since females can have heightened sensitivity to stress hormones compared to males (Bangasser and Wicks 2017; Daviu et al. 2014). Stress hormones are important modulators of memory generalization, and can increase the intensity or accelerate the natural time-course of memory generalization (Dos Santos Corrêa et al. 2019; Pedraza et al. 2016). Previous unpublished results from our lab compared adult females who had undergone several days of previous handling with those who simply remained in their homecages. Handled rats displayed significantly less fear when tested in the contextual fear conditioning 2 days after training compared to nonhandled subjects (**Figure 4.B**), despite not differing in anxiety behavior in the Elevated Plus-Maze (**Figure 4.C**), suggesting that frequent handling increased resilience to fear in these females, while having no effect on anxiety.

**Figure 4.**
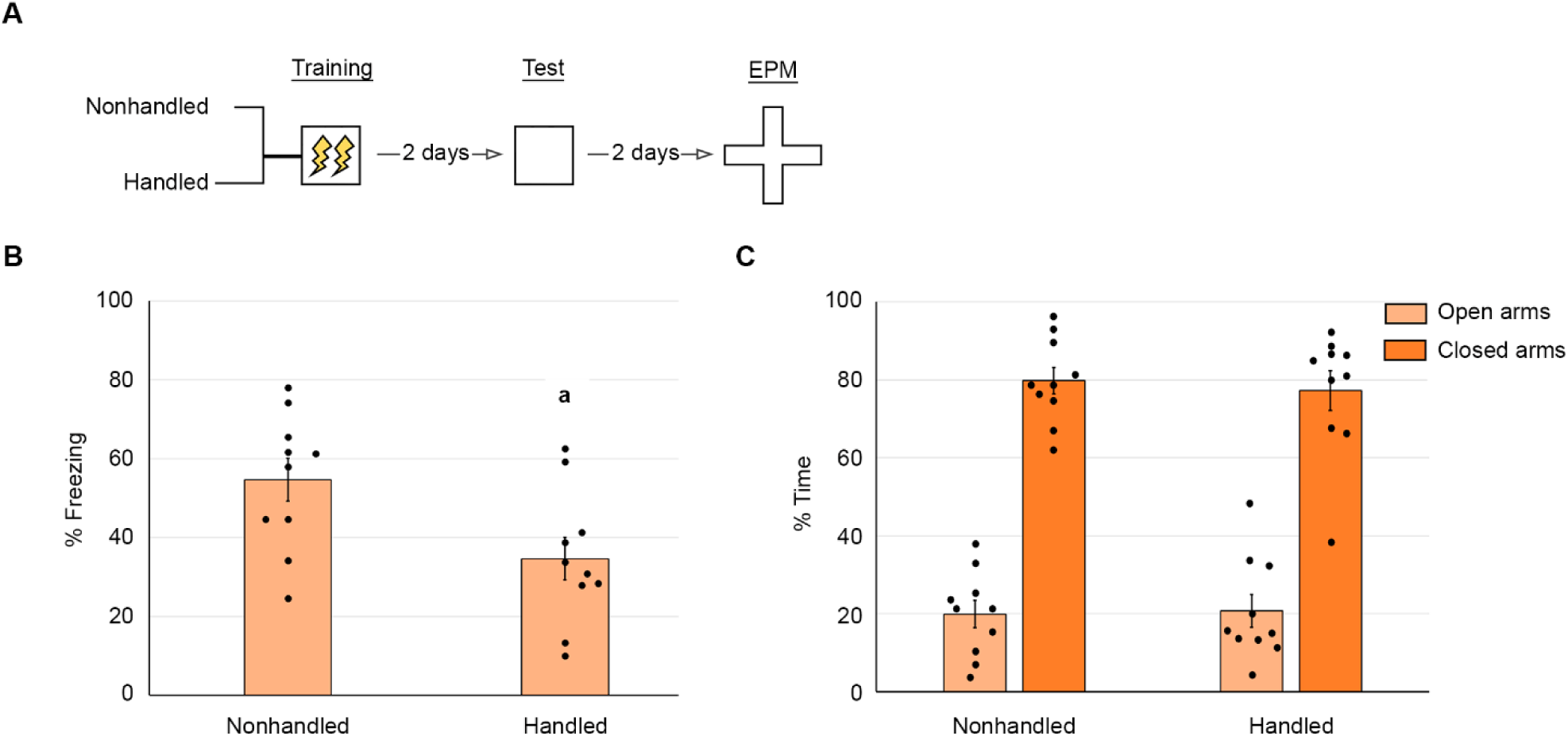
Comparison of conditioned fear and anxiety behavior in previously handled vs. nonhandled young adult female rats. (A) Experimental protocol. (B) Histogram showing mean±SEM of % freezing, with individual dots overlayed representing individual animals. Data analyzed by unpaired bicaudal Student’s t-test (t _(18)_ = 2.62; P < 0.0174). **a**: (different from nonhandled females). (C) Histogram showing mean±SEM of the % time spent in open and closed arms, with individual dots overlayed representing individual animals. Data analyzed by independent one-way ANOVA followed by Tukey *post hoc* test (F _(3,36)_ = 66.86, P < 0.0001). No significant difference was found between the mean time spent in the open and closed arms for handled and nonhandled rats.

In the CFC task, metal grid floors may be threat predictors, becoming more salient with time and facilitating fear generalization to neutral contexts (Huckleberry et al. 2016). Therefore, when testing a novel context D with a grid floor very similar to the one in the conditioning context A, males displayed significantly less fear in context D at 2 days (average 23% of session duration spent freezing), and a good increase at 28 and 45 days, a clear time-dependent generalization for this context (average 52% of session time at 45 days - **Figure S-2.B**). Conversely, females displayed similar freezing levels to males in context D at 2 days (average 22% of session time spent freezing) but showed less fear than their male counterparts when tested at 45 days (average 31% of session time - **Figure S-2.A**), even though freezing levels between males and females in the conditioning context did not differ significantly at this remote timepoint, indicating both sexes learned and retrieved the task equally well (**Figure S-2.C**). Taken together, these results suggest that while this salient aversive contextual cue (the grid) became progressively generalized to a novel context, other background features signaling safety for males might have become less salient, and counteracted the fear response, while the female group on average still perceived the context in its entirety as relatively safe.

There are pitfalls in comparing fear responses, between males and females based solely on observable behavioral differences. The CFC task is an inescapable, classical conditioning task, and even in auditory (tone) fear conditioning, subjects cannot escape the aversive context and may tend to express a more passive response, such as freezing in response to threat, rather than engaging in more active escape mechanisms such as flight or fight (Trott et al. 2022a) For instance, it is known that females are more prone to *darting*, a common defensive behavior in the presence of auditory cues: thus, freezing itself might not always be the taken as the most reliable quantifier of fear memory retention (Gruene et al. 2015). Furthermore, females can display heightened shock reactivity (Olivera-Pasilio and Dabrowska 2023) or distinct startle responses compared to males, although results investigating these differences have been mixed (Graham et al. 2009; Russo and Parsons 2021; Zhao et al. 2018). In our contextual fear conditioning experiments, neither males nor females were observed to engage in darting, and since most of our female subjects displayed relatively high levels of freezing in the conditioning context, we discarded the possibility of this defensive behavior interfering in our results. Interestingly, despite infrequent, males, but not females, displayed some *flight behavior* (jumping), depending on the context (data shown here): females in A (zero of 10 subjects), B (zero of 10), C (zero of 10) and D (zero of 11); males in A (1 jumper among 11 subjects), B (zero of 11), C (1 jumper among 11 subjects) and D (zero of 11).

### A CLOSER LOOK AT THE CHOSEN CONTEXTS: PCA

The main premise behind our selection of four graded contexts was that a “context” is, indeed, a *space-confined multisensory cluster* which involves multiple, inseparable (yet, distinguishable) parameters. Therefore, it is unreasonable to expect that controlling just one or two parameters at once in a pure contextual fear conditioning task is actually possible: animals integrate multiple sensory modalities to explore their environment, monitoring all modalities simultaneously, even when we attempt to isolate those where white noise masking is applicable (e.g., background sound and/or odor).

As previously described, our four contexts A, B, C and D were designed primarily on intuition, with contexts sorted according to multisensory criteria (tactile, visual, olfactory, or their combinations), which does not facilitate pinpointing which parameter, if any, dominates over the others for each sex, at each time interval. **Figure S-1** outlines the five chosen parameters – FLOOR, WALLS, SIZE, ROOM and SCENT, which, if varied simultaneously with each limited to just two possibilities (variables), would result in an unreasonably large number of combinations to test. Therefore, we reassessed our data by pooling groups according to the parameters and their variables to analyze correlations and perform multidimensional data reduction with Principal Component Analysis. The variables were, respectively, smooth vs. grid (FLOOR texture), planar vs. curved (WALL geometry), ethanol vs. isopropyl alcohol vs. quaternary ammonium (cleaning agent SCENT), large vs. small (chamber SIZE) and conditioning vs. novel (test session ROOM). Results are shown in **Figures S-3 B** and **C,** and **S-4 A, C** and **E**. For methodology of data regrouping and a detailed description of results see SUPPL. MATERIAL.

As illustrated in the heatmaps in **Figure S-3**, most of the measured behavioral responses (freezing levels) in each context were highly correlated, a condition for which Principal Components Analysis (PCA) is the ideal method for extracting useful information (Kirkwood et al. 2013).

**Figure S-4 A-F** shows the Principal Component Analysis results. No variables were fully negatively correlated (loadings at 180°), i.e., variables that exhibit different patterns of behavior, only positively correlated (clustered) ones - meaning they behave similarly – and orthogonal (loadings at 90°), i.e., uncorrelated variables – indicating independent variation.

Comparison of the four contexts A, B, C and D (**Figure S-4 A**) shows that while contexts A and B are always positively correlated in females, at 45 days this pattern changes for males, consistent with what is shown in **Figure 2**. When examining the covariation of all the 11 variants of the 5 parameters grouped collectively (**Figure S-4 C**), the most noticeable aspect is the clustering, at 2 days, of some of the variables that describe the conditioning (training) context A (grid floor, planar walls, small chamber and conditioning room, with the exception of SCENT), and their complete regrouping as an “A” cluster at 28 days, and specially for males at 45 days, suggesting a reinforcing role of conditioning variables for the retrieval of the aversive memory.

In **Figure S-4.E**, we explored the variables separated by parameter to distinguish sex-related and/or time-dependent differences, and the results were slightly different from those in **Figure S-4 C**, since the covariants differed. A certain degree of sexual dimorphism was observed in the independent (orthogonal) response of females and males to FLOOR texture (through tactile sense) at 2 days, however, floor texture was not a significant parameter in distinguishing between contexts in the recent memory test, although males tend to become better discriminators for this parameter with time compared to females. In relation to chamber SIZE, females distinguished the larger ones independently from males at 2 and 28 days, with males exhibiting this characteristic at 45 days. Neither sex distinguished the variations for the parameters WALL geometry, ROOM and Ethanol at 2 and 28 days, but these patterns were disrupted at 45 days, with males diverging in their responses and females converged on a response that did not effectively distinguish the variables.

In sum, multidimensional data reduction with Principal Component Analysis showed that both sexes did not discriminate between smooth vs. grid floors at first, but at more remote timepoints, the tactile perception of floor appeared to lose saliency for females, while males demonstrated a greater ability to distinguish between textures. Sexual dimorphism was also found for test chamber size, with females being more sensitive to this parameter at 2 and 28 days compared to males, but this sensitivity decreased over time. In contrast, males appeared to distinguish these variables better than females at 45 days: females seem to have poorer discriminatory ability with time for certain contextual features. No sexual dimorphisms were found for cleaning agent scent, wall geometry and test session room.

Despite the difficulty in separating parameters, multidimensional data reduction with Principal Component Analysis facilitated the identification of salient variables, showing that neither sex discriminated between smooth vs. grid floors at 2 days, however, tactile perception of floor texture appeared to gain saliency for males at more remote timepoints, when they began to distinguish between textures. Sexual dimorphism was also found for test chamber size, with females being more sensitive to this parameter at 2 and 28 days compared to males, but sensitivity decreases over time, and males exhibited better discrimination of chamber size than females at 45 days. In general, females appear to lose discriminatory ability over time for certain contextual features. No sexual differences were found for cleaning agent scent, wall geometry and – surprisingly – test session room, contradicting some previously published findings obtained with different experimental setups and in another strain or rats, Sprague-Dawley (Zhou and Riccio 1996).

### SEX-SPECIFIC COGNITIVE MAPS AND CONFIGURAL DISCRIMINATION

Our results suggest that different factors might be at stake in fear memory discrimination, such as distinct safety signaling perception between males and females, but with the generalization process occurring differently across sexes based on different context parameters. Dimorphic strategies have been observed in other hippocampus-dependent tasks, such as spatial navigation, in which circulating estrogen levels along with volumetric differences in the hippocampus and entorhinal cortex influenced the extent to which females adopt an allocentric in opposition to an egocentric navigation strategy (Keeley et al. 2013). In the case of auditory cued fear memories, contextual foreground cues are more salient in generating a fear response in females compared to males, while males in general use more background cues when retrieving contextual fear memories (Cossio et al. 2016), suggesting increased recruitment of the amygdala in females, while males may rely more on hippocampus-dependent processes. Naming a parameter in a contextual fear conditioning task a “cue” emphasizes a presumed salience for the subjects, which differs from “cued fear conditioning”, a distinct behavioral task that also involves generalization over time (Pollack et al. 2018). A cued fear conditioning task has at least one controllable parameter that strongly signals the recruitment of the expected behavioral response, such as a tone or light, thus being a simpler experimental setup compared to *pure* contextual fear conditioning, in which no single parameter is intentionally more prominent over the others. As discussed in the previous section, even in such a complex setup, some parameters may hold particular saliency, potentially due to an innate predisposition, and it is reasonable to question whether any of these intrinsically interconnected parameters -- as in the so-called configural view (Pearce and Bouton 2001) might serve as a cue that facilitates generalization. To discriminate or generalize fear-related contexts, subjects likely evaluate multiple environmental variables present in neutral contexts but may gradually assign differential weights to these features, relying on a Bayesian-like, probabilistic inference process to guide behavior (Valone 2024; Sanders et al. 2020).

While in our main experiment females expressed lower freezing levels than males in the conditioning context A at 2 days, groups tested at more remote timepoints equaled males in terms of freezing behavior, indicating *there was no deficit in acquisition or consolidation of the fear conditioning experience*, but what was in fact observed were differences in the qualitative transformation of long-term memory, consistent with numerous other studies (Taxier et al. 2020).

Another possibility to explain our findings may be that, at least in part, females and males have different *cognitive contextual maps* (Posani et al. 2018). While females might make use of other parameters to determine how safe or threatening a context is in its entirety, resulting in better context discrimination, relevant features, such as chamber size in recent memory, and floor texture at more remote intervals, appear to be two salient parameters that might predict to some extent the behavioral outcome.

At this point, it is again relevant to cite Roger Shepard (1987) (the italics are ours): “Because any object or situation experienced by an individual is *unlikely to recur in exactly the same form and context* (…) a full characterization of the change that even a single environmental event induces in an individual must entail a specification of how that individual’s behavioral dispositions have been altered relative to any ensuing situation. Differences in the way individuals of different species represent the same physical situation implicate, in each individual, an internal metric of similarity between possible situations”. At least in relation to rat behavior, we can say exactly the same for conspecifics of each sex.

## MATERIAL AND METHODS

### Subjects

Naïve male and female Wistar rats weighing 270–340g and aged 70–75 days at the beginning of experimental procedures, were used. All animals were provided by our local breeding colony (CREAL/UFRGS) and housed in plastic cages up to four individuals per cage. Males and females were housed with same-sex littermates at opposite ends of the same facility room, which was kept under a 12h light/dark cycle, controlled temperature of 21 ± 2°C and approximately 65% humidity. Experimental procedures were conducted during the light phase. Regular chow and water were available *ad libitum* for all animals. To prevent additional or differential stress, estrus cycle was not verified. All experiments were conducted in accordance to local animal care legislation and guidelines (Brazilian Federal Law 11,794/2008) and approved by the Committee on Ethics in the Use of Experimental Animals of the Federal University of Rio Grande do Sul (CEUA – UFRGS, Project # 40,945).

### Conditioning apparatus and novel contexts

All seven distinct contexts used in our pilot study and main experiment (**Figure S-1**) are described below.

The Conditioning Context A was the same for both experiments, and consisted of an internally illuminated Plexiglass box (L x W x H: 22 x 22 x 33 cm) with three black and one light brown wall. The floor was a grid of parallel 0.1cm caliber stainless steel bars spaced 1 cm apart. Rats were observed through a webcam mounted on the opaque upper lid. The conditioning chamber A was cleaned with 70% ethanol solution before placing each animal.

In the pilot study, Context P1 was, in our view, the least similar to Context A, and consisted of polyvinyl chloride (PVC) cylindric container (diameter: 25 cm; height: 22 cm) with vertical black and white stripes on its internal walls and a heavy transparent glass lid. The floor consisted of a smooth black plywood surface covered with wood shavings, which was replaced before placing each animal. The test chamber was cleaned with 70% isopropyl alcohol. A slightly modified version P1’, identical to P1 but without wood shavings, was also assayed.

Context P2 consisted of a scalene triangular chamber, smaller than Context C (sides x H: 35, 14.5, 23 x 6.5 cm), with brown plywood walls, white tempered glass floor and no lid. Context P2 was cleaned with quaternary ammonium disinfectant cleaning solution before each animal.

Context P3 was a modified, half-sized version of Context D (L x W x H: 25 x 25 x 29 cm), with dimensions similar to the Conditioning chamber A, and was cleaned with 70% isopropyl alcohol.

In our main experiment, Context B was a not-so-slightly modified version of Context A, in which the metal grid floor was covered with a smooth brown vinyl mat and a semi-circular rubber wall placed inside the conditioning chamber. Both Contexts A and B were cleaned with 70% ethanol before each session.

Context C consisted of a right triangular chamber (sides x H: 34, 28, 48 x 19 cm) with smooth black plywood floor and walls. Differently from A and B, Context C was cleaned with quaternary ammonium disinfectant prior to placing each animal.

Context D was a larger rectangular box (L x W x H: 50 x 25 x 29 cm) was placed in a different area of the conditioning room and contained three opaque beige walls, one transparent frontal wall and a semi-translucent lid. A desk lamp was placed beside the frontal wall. The floor consisted of a grid of parallel 0.1 cm caliber stainless copper bars spaced 1 cm apart. The test chamber was cleaned with 70% isopropyl alcohol.

Rats were kept in their home cages and habituated to the conditioning room under a constant fan background noise for at least 30 min before the start of each session. Contexts C, P1, P1’ and P2 were located in separate rooms from that of the conditioning chamber.

### Behavioral procedures

#### Contextual fear conditioning (CFC)

Rats were habituated to the conditioning context (A) for 3 min before receiving two 2-s continuous 0.5 mA footshocks separated by a 30-s interval. Thirty seconds after the second footshock rats were removed from the chamber and returned to their homecage. Males and females were not trained in the same experimental session.

#### Test sessions

For each training-test interval, conditioned animals were divided into four different groups, each one tested in a different test chamber: either A, P1/P1’, P2 or P3 – for the pilot study (**Figure 1**) or A, B, C or D - for the main experiment (**Figure 2**). In both experiments, test sessions lasted 5 min during which freezing responses were manually scored by an experienced observer using a stopwatch. Defensive freezing behavior, defined as the absence of all muscle movements except those related to breathing (Blanchard and Blanchard 1969), was used as an indicator of memory retention and expressed as a percentage of total session time. The number of attempted escapes (jumps) were also quantified as a secondary indicator of flight behavior. To prevent any eventual reactivation/reconsolidation of fear memory, which could influence the results, each rat group was tested in one context only. Contexts were not counterbalanced since this methodological approach reactivates memory and produces differences in fear expression based on test order (Huckleberry et al. 2016), and that this can be expressed differently in males and females (Keiser et al. 2017) adding an additional variable to our results. Males and females were never tested in the same experimental session.

### Statistical Analyses

After assessing data distribution for normality (Shapiro-Wilk test) and homoscedasticity (Levene’s test), we employed, for parametric data, the following tests to find statistical differences between groups (considering P < 0.05): for two independent groups, the unpaired two-tailed Student’s t-test was used, and for K groups, we employed independent one or two-way ANOVAs, followed by Tukey Multiple Comparisons *post hoc* test in order to sort the significant differences found. In order to support *reproducibility*, for every statistically significant (P < 0.05) effect, Effect Sizes were calculated, either using Hedge’s g (effect: small = 0.20, medium = 0.50, large = 0.80), or omega-squared (ω^2^), for the ANOVAs (effect: small = 0.02, medium = 0.06, large = 0.14). Outliers beyond two standard deviations above or below the mean were excluded from analyses.

In order to estimate the degree of discrimination between contexts, individual and average Z-scores for every neutral context were calculated and plotted: Z-scores inform where each individual score is located inside the normal curve in terms of standard deviations, compared to the average percentual freezing score for same sex control groups with the same training-test interval. In **Figures 1.D** and **2.H-J**, each point represents the behavioral response of one individual animal.

Pearson’s correlation analysis was used to compare mean freezing variables between different sexes, contextual features and training-test intervals and obtain p-values, correlation coefficient (r) and confidence intervals (CIs). Subsequently, Principal Component Analysis (PCA) was used to reduce multidimensional data into principal component (PC) variables which allow us to observe associations, identify patterns and verify outliers within multivariate data. Figures S-3 and S-4 summarize these findings. Pearson’s r coefficients, p-values and Confidence Intervals are shown in Table S-3. PCA loading scores, Eigenvalues and Explained Variance are displayed in Table S-4.

## ACKNOWLEDGMENTS

The authors kindly thank Isabel Cristina Marques Scarello for her invaluable technical assistance. We also thank Jociane de Carvalho Myskiw and Giuliana Petiz Zugno for helping to perform some of the experiments. Special thanks to João Paulo Maires Hoppe (Douglas Research Centre, McGill University) for his insightful comments on our multivariate analysis. This work was supported by fellowships and grants from the Conselho Nacional de Desenvolvimento Científico e Tecnológico (CNPq/Brazil), Coordenação de Aperfeiçoamento de Pessoal de Nível Superior (CAPES/Brazil), and Fundação de Apoio à Pesquisa do Rio Grande do Sul (FAPERGS/Brazil).

