## Supplemental Material for "Multiple context discrimination in adult rats: sex variability and dynamics of time-dependent generalization of an aversive memory"

#### INDEX

**Figure S-1** – Context configurations

***Comparing fear responses in each context***

**Figure S-2** - Comparing fear responses in each context

**Table S-1** - Significances and size effects of groups compared in Figure S-2

*Multivariate Analysis of behavioral data*

**Figure S-3** – Pearson's correlations - Heatmaps

**Table S-2** - Pearson's r coefficients, p-values and Confidence Intervals of groups analyzed in Figure S-3

**Figure S-4** - PCAs (A to F)

**Table S-3** - PCA loading scores, Eigenvalues and Explained Variances for groups analyzed in Figure S-4

#### *Context configurations*

Our initial goal was to obtain a graded fear response in the novel contexts in male and female rats, “stair-case like” representation of decreasing freezing levels at 2 days after training. The three neutral contexts B, C and D were designed primarily on intuition, to differ from the aversive conditioning chamber A in a graded form. The multiple novel contexts were intuitively conceived based on multidimensional criteria, namely chamber shape and floor texture (Pilot Study) and chamber size and odor (Main Experiment). **Table S-1**, below, illustrates the two sets of contexts and describe some of their distinguishing characteristics.

**A**

| Pilot Study | | | | |
|---|---|---|---|---|
| Context | 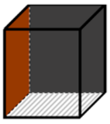 **A (conditioning)** | 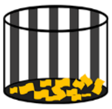 **P1 (neutral)** | 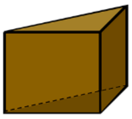 **P2 (neutral)** | 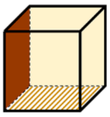 **P3 (neutral)** |
| Chamber shape | square cuboid | circular | triangular | square cuboid |
| Size | 22 x 22 x 33 (LxWxH cm) | 25 x 22 (D x H cm) | 35; 14.5; 23 x 6.5 (S1;S2;S3 x H cm) | 25 x 25 x 29 (LxWxH cm) |
| Floor texture | parellel 0.1 cm caliber stainless steel bars spaced 1 cm apart | smooth black plywood with wood shavings | white tempered glass | parallel 0.1 cm caliber stainless copper bars spaced 1 cm apart |
| Wall texture and color | three black and one pastel brown | vertical black and white stipes | brown plywood | three opaque beige and one transparent |
| Illumination | webcam and chamber wall lights | ceiling lamp | ceiling lamp | desk lamp near the transparent wall |
| Odor (cleaning agent) | 70% ethanol | 70% isopropyl alcohol | quaternary ammonium disinfectant | 70% isopropyl alcohol |
| Room used | conditioning room | novel room | novel room | conditioning room |

**B**

| Main Experiment | | | | |
|---|---|---|---|---|
| Context | 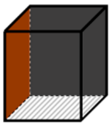 **A (conditioning)** | 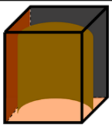 **B (modified conditioning)** | 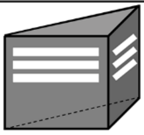 **C (neutral)** | 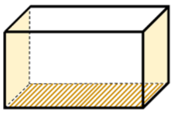 **D (neutral)** |
| Chamber shape | square cuboid | semicircle | triangular | rectangular cuboid |
| Size | 22 x 22 x 33 (LxWxH cm) | 25 x 22 x 33 (LxWxH cm) | 34; 28; 48 x 19 (S1;S2;S3 x H cm) | 50 x 25 x 29 (LxWxH cm) |
| Floor texture | parellel 0.1 cm caliber stainless steel bars spaced 1 cm apart | smooth brown vinyl | smooth black plywood | parallel 0.1 cm caliber stainless copper bars spaced 1 cm apart |
| Wall texture and color | three black and one pastel brown | same as context A with an added brown rubber forming a semicircle | smooth black plywood with white markings | three opaque beige and one transparent |
| Illumination | webcam and chamber wall lights | webcam light | ceiling lamp | desk lamp near the transparent wall |
| Odor (cleaning agent) | 70% ethanol | 70% ethanol | quaternary ammonium disinfectant | 70% isopropyl alcohol |
| Room used | conditioning room | conditioning room | novel room | conditioning room |

**Supplemental Figure 1.** schematic illustration depicting the specific contexts used in this study. (**A**) Pilot study (2 days only); (**B**) Main experiment (2, 28 and 45 day training-test intervals). In our pilot groups we also included P1', a modified version of context P1, without wood shavings in the floor.

#### *Comparing fear responses in each context*

We compared the performance of females and males separately across these time intervals – in contexts A, B, C and D. The results are presented in the Supplemental **Figure S-2.**, below.

##### Context A:

Independent two-way ANOVA of rats tested in context A in three different training-test intervals revealed only the effect of Sex ($F_{(1,55)} = 25.13$; $P < 0.0001$, $\omega2 = 0.28$), with no significant effect either in the interaction Sex x Time ($F_{(2,55)} = 2.389$; $P = 0.1012$), or in Time ($F_{(2,55)} = 2.403$; $P = 0.0999$). According to the ω2, the detected effect was large and robust.

In females, Tukey *post hoc* multiple comparisons revealed a negligible difference in freezing levels for context A between 2 and 28 days ($P = 0.0422$, g =1.557), and no significant differences between 28 and 45 days ($P = 0.6499$). In males, there were no significant differences between 2 and 28 days ($P > 0.999$) and between 28 and 45 days ($P = 0.9983$). Comparing sexes, females differed from males only at 2 days ($P = 0.0004$, g = 2.70), while no significant differences were found at 28 ($P = 0.6029$), and 45 days ($P = 0.1418$). Number of subjects per sex at each time point: females (2, 28 and 45d) = 10, 11, 9, and males = 11, 11, 9.

**Context B:**

For animals tested in the modified conditioning context B, independent two-way ANOVA revealed an effect of Sex ($F_{(1,57)} = 45.58$; $P < 0.0001$, ω2 = 0.41), Time ($F_{(2,57)} = 29.72$; $P < 0.0001$, ω2 = 0.48) and a Sex x Time interaction ($F_{(2,57)} = 11.53$; $P < 0.001$, ω2 = 0.25). According to the ω2, the detected effect was large and robust.

In females, Tukey *post hoc* multiple comparisons revealed a significant difference in freezing levels 2 and 28 days ($P < 0.0001$, g =4.229), and no significant differences between 28 and 45 days ($P = 0.1045$). In males, there were no significant differences between 2 and 28 days ($P = 0.1823$) and between 28 and 45 days ($P = 0.2479$). Comparing sexes, females differed from males only at 2 days ($P < 0.0001$, g = 5.320), while no significant differences were found at 28 ($P = 0.5256$), and 45 days ($P = 0.2679$). Number of subjects per sex at each time point: females (2, 28 and 45d) = 10, 11, 9, and males = 11, 11, 11.

**Context C:**

For animals tested in the novel context C, independent two-way ANOVA revealed an effect of Sex ($F_{(1,58)} = 45.58$; $P < 0.0001$, ω2 = 0.41), but no effect of Time ($F_{(2,58)} = 2.165$; $P = 0.1239$), or Sex x Time interaction ($F_{(1,58)} = 1.815$; $P = 0.172$). According to the ω2, the detected effect was large and robust.

In females, Tukey *post hoc* multiple comparisons revealed there were no differences in freezing levels 2 and 28 days ($P = 0.0926$), or between 28 and 45 days (P = 0.9064). Likewise in males, no differences were found between groups tested at 2 and 28 days (P >0.9999) and between 28 and 45 days (P = 0.9834). Comparing sexes, females differed from males at 2 days (P = 0.0009, g = 2.909), and 45 days (P = 0.0175, g = 1.199), but not at 28 days (P = 0.4641). Number of subjects per sex at each time point: females (2, 28 and 45d) = 10, 12, 9, and males = 11, 11, 11.

**Context D:**

For rats tested in the novel context D, independent two-way ANOVA revealed an effect of Sex ($F_{(1,58)} = 4.651$; P = 0.2571, ω2 = 0.05), and of Time ($F_{(2,58)} = 6.831$; P = 0.0022, ω2 = 0.15), but not a Sex x Time interaction ($F_{(2,58)} = 1.391$; P = 0.2571). According to the ω2, the detected effect for Time was large and robust.

In females, Tukey *post hoc* multiple comparisons revealed there were no differences in freezing levels 2 and 28 days (P = 0.4755), or between 28 and 45 days (P = 0.9893). In males, no differences were found between groups tested at 2 and 28 days (P = 0.1024) and between 28 and 45 days (P = 0.9453), but there was a significant difference between males tested at 2 and 45 days (P = 0.0097, g = 1.895). Comparing sexes, females did not differ from males at any of the tested time intervals (P > 0.15 in all cases). Number of subjects per sex at each time point: females (2, 28 and 45d) = 11, 11, 9, and males = 11, 11, 11.

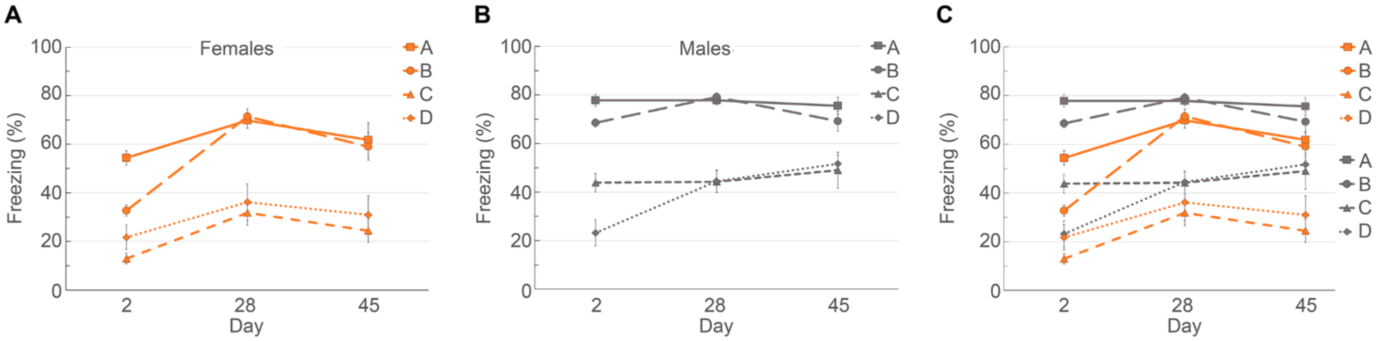


**Supplemental Figure 2.** comparisons of performances for each sex and timepoint in each context**.** (A) Line graphs representing freezing responses of females tested at 2, 28 and 45 days in contexts A, B, C and D. (B) *idem* for males tested at the same time points in the same contexts. (C) Comparison between males and females.

The comparisons shown above between male and female rats in each context tested in the Pilot Study and Main Experiment, were based on the adjusted p values (independent 2-way ANOVA with Tukey's posthoc test) presented in **Table S-1**, below, which includes Hedge's g calculated for cases in which P > 0.05.

| **Pilot Experiment** | | |
|---|---|---|
| **Tukey's multiple comparisons test** | **Effect size (g)** | **Adjusted P Value** |
| Females:Context A vs. Females:Context P1 | 1.896 | <0.0001 |
| Females:Context A vs. Females:Context P2 | 4.55 | <0.0001 |
| Females:Context A vs. Females:Context P3 | 3.848 | <0.0001 |
| Females:Context A vs. Females:Context P1' | 2.069 | 0.0001 |
| Females:Context A vs. Males:Context A | - | >0.9999 |
| Females:Context A vs. Males:Context P1 | - | >0.9999 |
| Females:Context A vs. Males:Context P2 | 2.44 | 0.0083 |
| Females:Context A vs. Males:Context P3 | 2.994 | 0.014 |
| Females:Context A vs. Males:Context P1' | - | 0.7236 |
| Females:Context P1 vs. Females:Context P2 | - | 0.9753 |
| Females:Context P1 vs. Females:Context P3 | - | >0.9999 |
| Females:Context P1 vs. Females:Context P1' | - | >0.9999 |
| Females:Context P1 vs. Males:Context A | 1.979 | <0.0001 |
| Females:Context P1 vs. Males:Context P1 | 1.656 | <0.0001 |
| Females:Context P1 vs. Males:Context P2 | 0.483 | 0.8786 |
| Females:Context P1 vs. Males:Context P3 | 0.523 | 0.8648 |
| Females:Context P1 vs. Males:Context P1' | 1.133 | 0.0445 |
| Females:Context P2 vs. Females:Context P3 | - | >0.9999 |
| Females:Context P2 vs. Females:Context P1' | - | 0.9646 |
| Females:Context P2 vs. Males:Context A | 4.473 | <0.0001 |
| Females:Context P2 vs. Males:Context P1 | 3.128 | <0.0001 |
| Females:Context P2 vs. Males:Context P2 | - | 0.324 |
| Females:Context P2 vs. Males:Context P3 | - | 0.3182 |
| Females:Context P2 vs. Males:Context P1' | 2.901 | 0.0056 |
| Females:Context P3 vs. Females:Context P1' | - | 0.9996 |
| Females:Context P3 vs. Males:Context A | 3.833 | <0.0001 |
| Females:Context P3 vs. Males:Context P1 | 2.725 | <0.0001 |
| Females:Context P3 vs. Males:Context P2 | - | 0.691 |
| Females:Context P3 vs. Males:Context P3 | - | 0.675 |
| Females:Context P3 vs. Males:Context P1' | 2.353 | 0.0278 |
| Females:Context P1' vs. Males:Context A | 2.156 | <0.0001 |
| Females:Context P1' vs. Males:Context P1 | 1.737 | 0.0003 |
| Females:Context P1' vs. Males:Context P2 | - | 0.9664 |
| Females:Context P1' vs. Males:Context P3 | - | 0.9586 |
| Females:Context P1' vs. Males:Context P1' | - | 0.1204 |

| | | |
|---|---|---|
| Males:Context A vs. Males:Context P1 | - | 0.9996 |
| Males:Context A vs. Males:Context P2 | 2.49 | 0.0029 |
| Males:Context A vs. Males:Context P3 | 2.962 | 0.0054 |
| Males:Context A vs. Males:Context P1' | - | 0.578 |
| Males:Context P1 vs. Males:Context P2 | 1.727 | 0.0189 |
| Males:Context P1 vs. Males:Context P3 | 1.872 | 0.0313 |
| Males:Context P1 vs. Males:Context P1' | - | 0.904 |
| Males:Context P2 vs. Males:Context P3 | - | >0.9999 |
| Males:Context P2 vs. Males:Context P1' | - | 0.7376 |
| Males:Context P3 vs. Males:Context P1' | - | 0.8 |
| **Main Experiment** | | |
| **Tukey's multiple comparisons test** | **Effect size (g)** | **Adjusted P Value** |
| **2 days** | | |
| Females:Context A vs. Females:Context B | 2.256 | 0.002 |
| Females:Context A vs. Females:Context C | 4.966 | <0.0001 |
| Females:Context A vs. Females:Context D | 2.379 | <0.0001 |
| Females:Context A vs. Males:Context A | 2.641 | 0.0004 |
| Females:Context A vs. Males:Context B | - | 0.1147 |
| Females:Context A vs. Males:Context C | - | 0.4441 |
| Females:Context A vs. Males:Context D | 2.16 | <0.0001 |
| Females:Context B vs. Females:Context C | 2.644 | 0.0068 |
| Females:Context B vs. Females:Context D | - | 0.3884 |
| Females:Context B vs. Males:Context A | 5.566 | <0.0001 |
| Females:Context B vs. Males:Context B | 5.294 | <0.0001 |
| Females:Context B vs. Males:Context C | - | 0.3677 |
| Females:Context B vs. Males:Context D | - | 0.5778 |
| Females:Context C vs. Females:Context D | - | 0.6724 |
| Females:Context C vs. Males:Context A | 8.247 | <0.0001 |
| Females:Context C vs. Males:Context B | 8.583 | <0.0001 |
| Females:Context C vs. Males:Context C | 2.933 | <0.0001 |
| Females:Context C vs. Males:Context D | - | 0.4789 |
| Females:Context D vs. Males:Context A | 4.24 | <0.0001 |
| Females:Context D vs. Males:Context B | 3.745 | <0.0001 |
| Females:Context D vs. Males:Context C | 1.486 | 0.0007 |
| Females:Context D vs. Males:Context D | - | >0.9999 |
| Males:Context A vs. Males:Context B | - | 0.5719 |
| Males:Context A vs. Males:Context C | 3.132 | <0.0001 |
| Males:Context A vs. Males:Context D | 3.923 | <0.0001 |
| Males:Context B vs. Males:Context C | 2.483 | 0.0001 |
| Males:Context B vs. Males:Context D | 3.426 | <0.0001 |
| Males:Context C vs. Males:Context D | 1.331 | 0.002 |
| **28 days** | | |
| Females:Context A vs. Females:Context B | - | >0.9999 |
| Females:Context A vs. Females:Context C | 2.54 | <0.0001 |
| Females:Context A vs. Females:Context D | 1.758 | <0.0001 |

| | | |
|---|---|---|
| Females:Context A vs. Males:Context A | - | 0.903 |
| Females:Context A vs. Males:Context B | - | 0.8021 |
| Females:Context A vs. Males:Context C | | 0.0023 |
| Females:Context A vs. Males:Context D | | 0.0026 |
| Females:Context B vs. Females:Context C | 2.647 | <0.0001 |
| Females:Context B vs. Females:Context D | 1.854 | <0.0001 |
| Females:Context B vs. Males:Context A | - | 0.9711 |
| Females:Context B vs. Males:Context B | - | 0.918 |
| Females:Context B vs. Males:Context C | 1.999 | 0.0009 |
| Females:Context B vs. Males:Context D | 1.905 | 0.001 |
| Females:Context C vs. Females:Context D | - | 0.9965 |
| Females:Context C vs. Males:Context A | 3.339 | <0.0001 |
| Females:Context C vs. Males:Context B | 3.479 | <0.0001 |
| Females:Context C vs. Males:Context C | - | 0.4628 |
| Females:Context C vs. Males:Context D | - | 0.4405 |
| Females:Context D vs. Males:Context A | 2.289 | <0.0001 |
| Females:Context D vs. Males:Context B | 2.381 | <0.0001 |
| Females:Context D vs. Males:Context C | - | 0.8957 |
| Females:Context D vs. Males:Context D | - | 0.8824 |
| Males:Context A vs. Males:Context B | - | >0.9999 |
| Males:Context A vs. Males:Context C | 2.967 | <0.0001 |
| Males:Context A vs. Males:Context D | 2.803 | <0.0001 |
| Males:Context B vs. Males:Context C | 3.145 | <0.0001 |
| Males:Context B vs. Males:Context D | 2.967 | <0.0001 |
| Males:Context C vs. Males:Context D | - | >0.9999 |
| **45 days** | | |
| Females:Context A vs. Females:Context B | - | >0.9999 |
| Females:Context A vs. Females:Context C | 2.063 | 0.0008 |
| Females:Context A vs. Females:Context D | 1.378 | 0.0105 |
| Females:Context A vs. Males:Context A | - | 0.725 |
| Females:Context A vs. Males:Context B | - | 0.9819 |
| Females:Context A vs. Males:Context C | - | 0.7521 |
| Females:Context A vs. Males:Context D | - | 0.9097 |
| Females:Context B vs. Females:Context C | 2.219 | 0.0025 |
| Females:Context B vs. Females:Context D | 1.377 | 0.0278 |
| Females:Context B vs. Males:Context A | - | 0.5093 |
| Females:Context B vs. Males:Context B | - | 0.904 |
| Females:Context B vs. Males:Context C | - | 0.9143 |
| Females:Context B vs. Males:Context D | - | 0.9836 |
| Females:Context C vs. Females:Context D | - | 0.9939 |
| Females:Context C vs. Males:Context A | 4.055 | <0.0001 |
| Females:Context C vs. Males:Context B | 3.227 | <0.0001 |
| Females:Context C vs. Males:Context C | - | 0.0578 |
| Females:Context C vs. Males:Context D | 2.046 | 0.0233 |
| Females:Context D vs. Males:Context A | 2.45 | <0.0001 |

| | | |
|---|---|---|
| Females:Context D vs. Males:Context B | 2.058 | 0.0002 |
| Females:Context D vs. Males:Context C | - | 0.3329 |
| Females:Context D vs. Males:Context D | - | 0.1779 |
| Males:Context A vs. Males:Context B | - | 0.9932 |
| Males:Context A vs. Males:Context C | 1.348 | 0.0299 |
| Males:Context A vs. Males:Context D | - | 0.0723 |
| Males:Context B vs. Males:Context C | - | 0.1519 |
| Males:Context B vs. Males:Context D | - | 0.3023 |
| Males:Context C vs. Males:Context D | - | >0.9999 |

**Supplemental Table 1.** P values obtained in the Pilot Study and Main Experiment (independent two-way ANOVA with Tukey posthoc test). Hedge's g effect sizes were added where applicable.

#### *Multivariate Analysis of behavioral data*

##### Conceptualization and methodology

It remains unclear which *boundary conditions* mediate the selection between slightly different contexts. However, we adhere to the configural perspective (Pearce and Bouton, 2001), which posits that a whole set of sensory perceptions becomes associated simultaneously with an unconditioned stimulus, i.e., a 'context' is understood as a *space-confined multisensory cluster* composed of inseparable yet distinguishable parameters.

Similar to the three Pilot neutral contexts, neutral contexts B, C, and D were designed intuitively to differ from the aversive conditioning chamber A in a graded manner. Each incorporated multiple types of sensory information (tactile, visual, olfactory, or combinations thereof) in order to obtain measurable differences, despite the difficulty in determining which modality might dominate for each sex at different times after training. **Figure S-1** outlines the several parameters from which five were selected – FLOOR, WALLS, SIZE, ROOM and SCENT. For Multivariate Analysis, two or three variables were attributed to each of these five parameters: respectively, smooth vs. grid (FLOOR texture), planar vs. curved (WALL geometry), ethanol vs. isopropyl alcohol vs. quaternary ammonium (cleaning agent SCENT), larger vs. smaller (chamber SIZE) and conditioning vs. novel (test session ROOM).

Each of the eleven groups described above was formed by *pooling* data from two or three context groups, selected based on the relevant parameter, for instance, data from contexts A + D for contexts with a grid floor, and from B + C for smooth floor. Groupings were as follows: A+D (grid) vs. B+C (smooth) for FLOOR, A+C+D (planar) vs. B (curved), for WALL, A+B (smaller) vs. C+D (larger) for SIZE, A+B+D (conditioning) vs. C (novel) for ROOM, and A+B (ethanol) vs. C (quaternary ammonium) vs. D (isopropyl alcohol). Note that with these five parameters, there were no redundant groupings - each pool was distinct. Because the pooled groups differed in sample size, missing values were replaced with the respective sample mean to perform analyses.

We then reassessed our reorganized data analyzing correlations (Heatmaps) and performing multidimensional data reduction with Principal Component Analysis. Results are shown in **Figures S-3** and **S-4.**

#### *Pearson's Correlation Matrices*

We performed Pearson's correlation analyses to identify positively and negatively correlated sensory modality variables between females and males tested at each post-training interval (2, 28 and 45 days). **Figure S-3** below shows the Heatmaps of (A) the four contexts A, B, C and D in the main experiment, organized per sex and training-test interval, (B) the 11 variables of the 5 selected parameters compared simultaneously - organized as in (A), and (C) the 5 parameters analyzed separately, comparing their respective variables per sex.

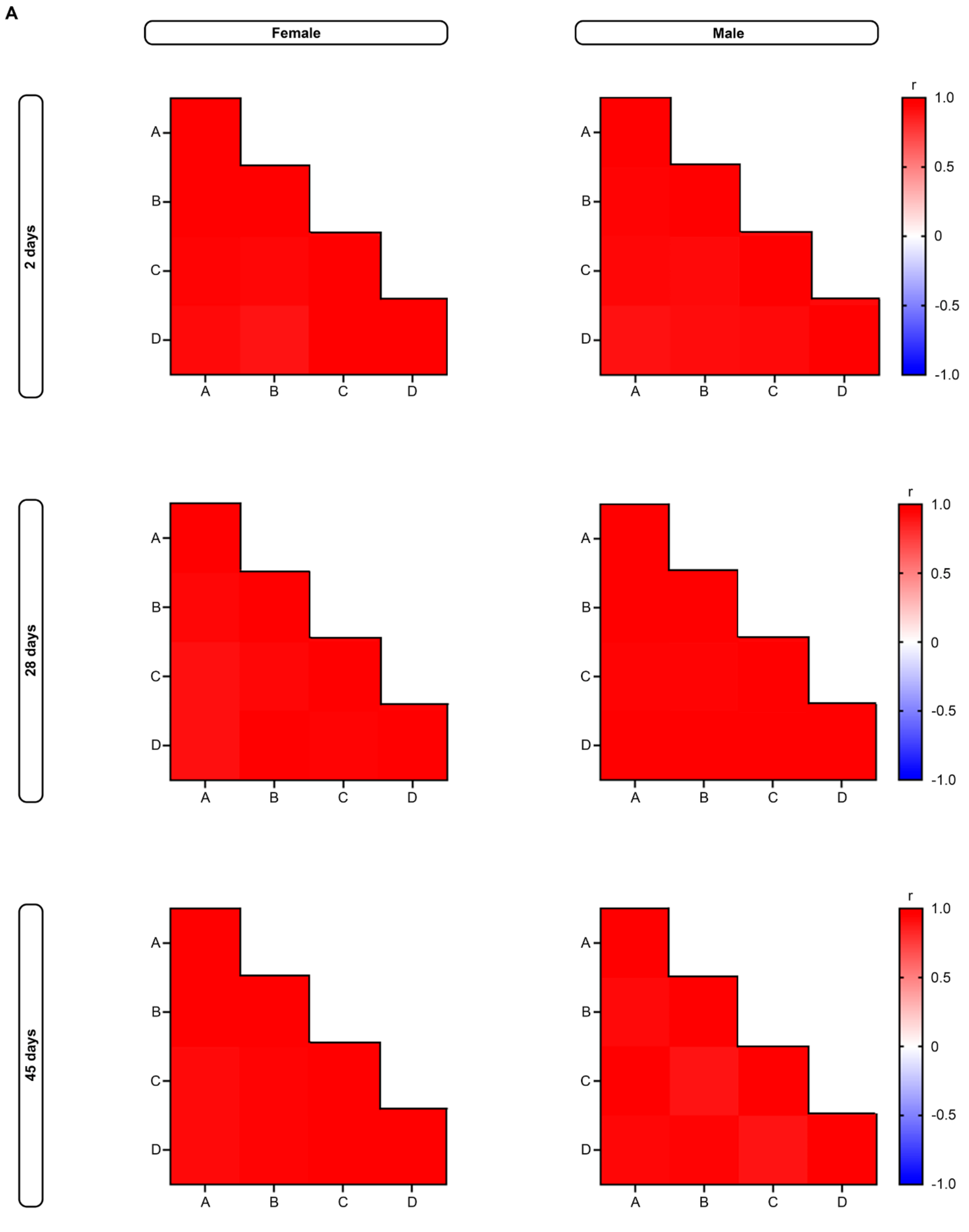

A
Female
Male
2 days
28 days
45 days
A
B
C
D
r
1.0
0.5
0
-0.5
-1.0

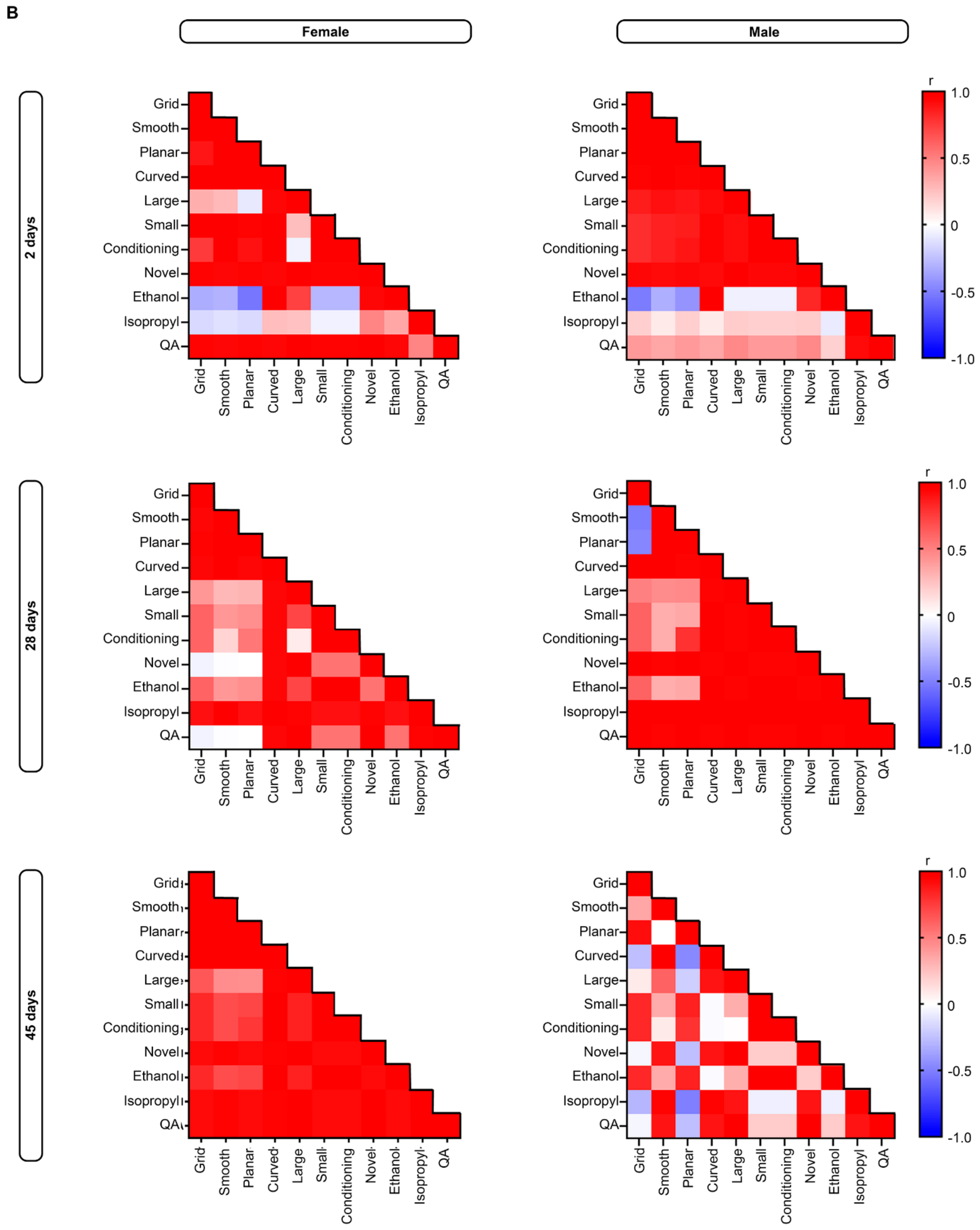
B
Female
Male
2 days
28 days
45 days
Grid
Smooth
Planar
Curved
Large
Small
Conditioning
Novel
Ethanol
Isopropyl
QA
r
1.0
0.5
0
-0.5
-1.0

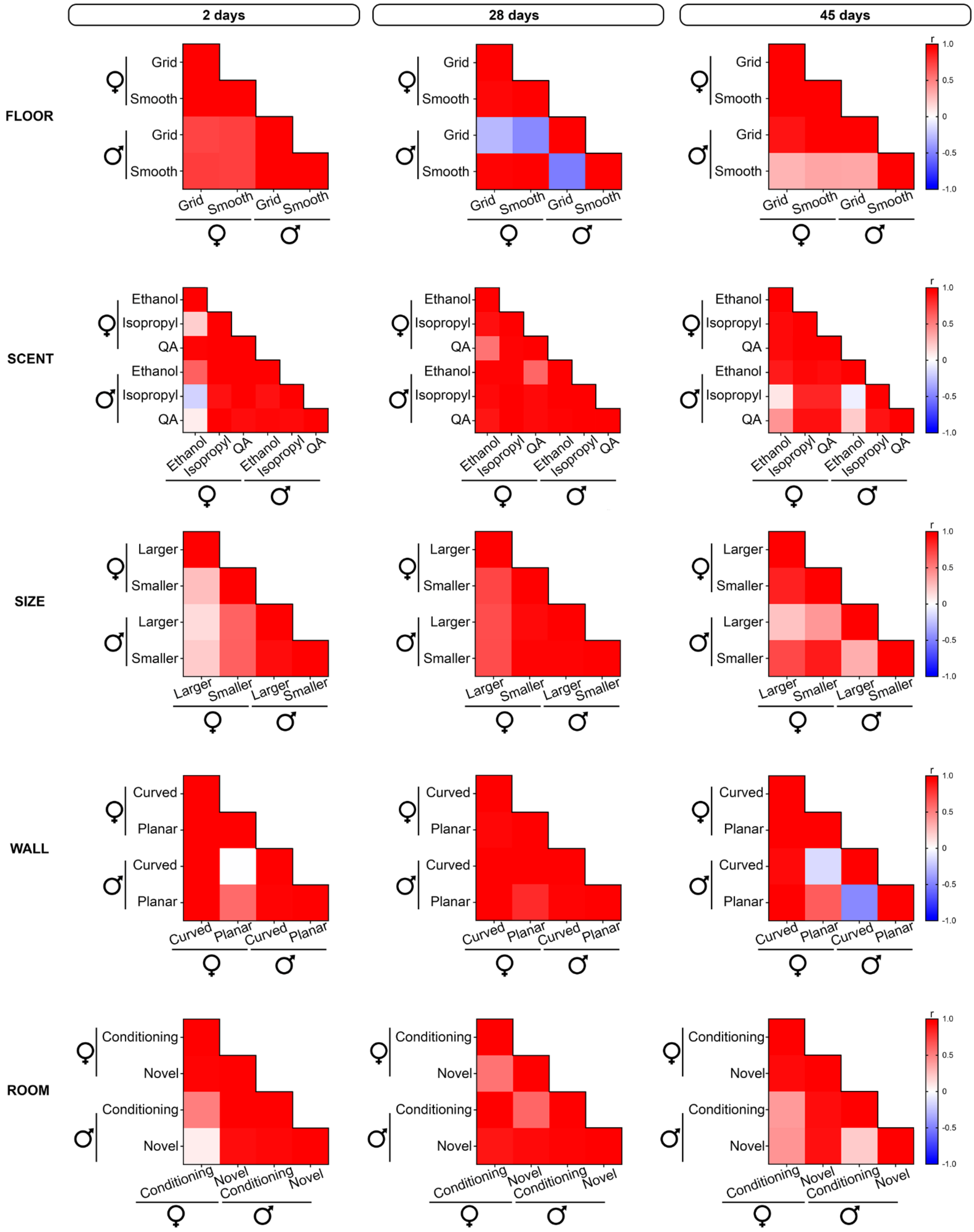


**Supplemental Figure 3.** Pearson's correlation matrix heatmaps for different contexts and parameter groups (Main Experiment). **A:** Pearson's correlation coefficients comparing mean percentage of freezing time in each test context (A, B, C and D) for females and males at 2, 28 and 45 days. **B:** Correlations for each of the eleven parameter variables (grid vs. smooth, curved vs. planar, larger vs. smaller, conditioning vs. novel, ethanol vs. isopropyl alcohol vs. quaternary ammonium) compared across sexes and different intervals. **C:** Heatmaps showing correlations between different sensory modality parameters (Cleaning agent scent, Wall geometry, Test session room, Floor texture and Test chamber size) for each interval. Square color shows the Pearson's r coefficient (r), with dark red indicating a strong positive correlation (r = +1) and dark blue, a strong negative correlation (r = -1). White squares indicate no correlation between comparisons (r = 0). Percentage of freezing times were pooled together for Smooth floor (contexts B and C), Grid floor (contexts A and D), Curved walls (context B) and Planar walls (contexts A, C and D), Large test chamber (contexts C and D) and Small test chamber (contexts A and B), Conditioning room (contexts A, B and D) and Novel room (context C), 70% ethanol (contexts A and B), 70% isopropyl (context D) and quaternary ammonium (context C). QA: quaternary ammonium.

### Correlation Matrices

| Pearson's r | | | | |
|---|---|---|---|---|
| **2 days** | | | | |
| | **Larger chamber females** | **Smaller chamber females** | **Larger chamber males** | **Smaller chamber males** |
| **Larger chamber females** | 1.000 | 0.260 | 0.144 | 0.208 |
| **Smaller chamber females** | 0.260 | 1.000 | 0.615 | 0.622 |
| **Larger chamber males** | 0.144 | 0.615 | 1.000 | 0.910 |
| **Smaller chamber males** | 0.208 | 0.622 | 0.910 | 1.000 |
| **28 days** | | | | |
| | **Larger chamber females** | **Smaller chamber females** | **Larger chamber males** | **Smaller chamber males** |
| **Larger chamber females** | 1.000 | 0.719 | 0.687 | 0.691 |
| **Smaller chamber females** | 0.719 | 1.000 | 0.925 | 0.948 |
| **Larger chamber males** | 0.687 | 0.925 | 1.000 | 0.950 |
| **Smaller chamber males** | 0.691 | 0.948 | 0.950 | 1.000 |
| **45 days** | | | | |
| | **Larger chamber females** | **Smaller chamber females** | **Larger chamber males** | **Smaller chamber males** |
| **Larger chamber females** | 1.000 | 0.849 | 0.245 | 0.710 |
| **Smaller chamber females** | 0.849 | 1.000 | 0.408 | 0.868 |
| **Larger chamber males** | 0.245 | 0.408 | 1.000 | 0.322 |
| **Smaller chamber males** | 0.710 | 0.868 | 0.322 | 1.000 |

| P-values | | | | |
|---|---|---|---|---|
| **2 days** | | | | |
| | **Larger chamber females** | **Smaller chamber females** | **Larger chamber males** | **Smaller chamber males** |
| **Larger chamber females** | 0 | 0.267947561 | 0.534463436 | 0.364648606 |
| **Smaller chamber females** | 0.267947561 | 0 | 0.003919033 | 0.003384902 |
| **Larger chamber males** | 0.534463436 | 0.003919033 | 0 | 4.25778E-09 |
| **Smaller chamber males** | 0.364648606 | 0.003384902 | 4.25778E-09 | 0 |
| **28 days** | | | | |
| | **Larger chamber females** | **Smaller chamber females** | **Larger chamber males** | **Smaller chamber males** |
| **Larger chamber females** | 0 | 0.000160709 | 0.000409887 | 0.000366955 |
| **Smaller chamber females** | 0.000160709 | 0 | 7.45724E-10 | 1.97506E-11 |
| **Larger chamber males** | 0.000409887 | 7.45724E-10 | 0 | 1.30209E-11 |
| **Smaller chamber males** | 0.000366955 | 1.97506E-11 | 1.30209E-11 | 0 |
| **45 days** | | | | |
| | **Larger chamber females** | **Smaller chamber females** | **Larger chamber males** | **Smaller chamber males** |
| **Larger chamber females** | 0 | 8.34521E-06 | 0.326721358 | 0.00097313 |
| **Smaller chamber females** | 8.34521E-06 | 0 | 0.092568749 | 2.99982E-06 |
| **Larger chamber males** | 0.326721358 | 0.092568749 | 0 | 0.166113178 |
| **Smaller chamber males** | 0.00097313 | 2.99982E-06 | 0.166113178 | 0 |

| Confidence Intervals of r | | | | |
|---|---|---|---|---|
| **2 days** | | | | |
| | **Larger chamber females** | **Smaller chamber females** | **Larger chamber males** | **Smaller chamber males** |
| **Larger chamber females** | 1.000 to 1.000 | -0.2061 to 0.6301 | -0.3071 to 0.5417 | -0.2454 to 0.5873 |
| **Smaller chamber females** | -0.2061 to 0.6301 | 1.000 to 1.000 | 0.2366 to 0.8312 | 0.2482 to 0.8349 |
| **Larger chamber males** | -0.3071 to 0.5417 | 0.2366 to 0.8312 | 1.000 to 1.000 | 0.7927 to 0.9624 |
| **Smaller chamber males** | -0.2454 to 0.5873 | 0.2482 to 0.8349 | 0.7927 to 0.9624 | 1.000 to 1.000 |
| **28 days** | | | | |
| | **Larger chamber females** | **Smaller chamber females** | **Larger chamber males** | **Smaller chamber males** |
| **Larger chamber females** | 1.000 to 1.000 | 0.4275 to 0.8755 | 0.3740 to 0.8597 | 0.3806 to 0.8617 |
| **Smaller chamber females** | 0.4275 to 0.8755 | 1.000 to 1.000 | 0.8252 to 0.9688 | 0.8776 to 0.9787 |
| **Larger chamber males** | 0.3740 to 0.8597 | 0.8252 to 0.9688 | 1.000 to 1.000 | 0.8825 to 0.9796 |
| **Smaller chamber males** | 0.3806 to 0.8617 | 0.8776 to 0.9787 | 0.8825 to 0.9796 | 1.000 to 1.000 |
| **45 days** | | | | |
| | **Larger chamber females** | **Smaller chamber females** | **Larger chamber males** | **Smaller chamber males** |
| **Larger chamber females** | 1.000 to 1.000 | 0.6332 to 0.9424 | -0.2503 to 0.6389 | 0.3629 to 0.8837 |
| **Smaller chamber females** | 0.6332 to 0.9424 | 1.000 to 1.000 | -0.07240 to 0.7350 | 0.6749 to 0.9500 |
| **Larger chamber males** | -0.2503 to 0.6389 | -0.07240 to 0.7350 | 1.000 to 1.000 | -0.1405 to 0.6692 |
| **Smaller chamber males** | 0.3629 to 0.8837 | 0.6749 to 0.9500 | -0.1405 to 0.6692 | 1.000 to 1.000 |

### Correlation Matrices

#### Pearson's r

##### 2 days

| | Conditioning room females | Novel room females | Conditioning room males | Novel room males |
|---|---|---|---|---|
| **Conditioning room females** | 1.00 | 0.95 | 0.51 | 0.07 |
| **Novel room females** | 0.95 | 1.00 | 0.97 | 0.92 |
| **Conditioning room males** | 0.51 | 0.97 | 1.00 | 0.93 |
| **Novel room males** | 0.07 | 0.92 | 0.93 | 1.00 |

##### 28 days

| | Conditioning room females | Novel room females | Conditioning room males | Novel room males |
|---|---|---|---|---|
| **Conditioning room females** | 1.00 | 0.55 | 0.96 | |
| **Novel room females** | 0.55 | 1.00 | 0.60 | 0.92 |
| **Conditioning room males** | 0.96 | 0.60 | 1.00 | 0.95 |
| **Novel room males** | 0.89 | 0.92 | 0.95 | 1.00 |

##### 45 days

| | Conditioning room females | Novel room females | Conditioning room males | Novel room males |
|---|---|---|---|---|
| **Conditioning room females** | 1.00 | 0.92 | 0.40 | 0.42 |
| **Novel room females** | 0.92 | 1.00 | 0.92 | 0.90 |
| **Conditioning room males** | 0.40 | 0.92 | 1.00 | 0.21 |
| **Novel room males** | 0.42 | 0.90 | 0.21 | 1.00 |

#### P-values

##### 2 days

| | Conditioning room females | Novel room females | Conditioning room males | Novel room males |
|---|---|---|---|---|
| **Conditioning room females** | 0 | 1.88803E-05 | 0.003457138 | 0.835480603 |
| **Novel room females** | 1.88803E-05 | 0 | 6.0366E-06 | 0.000192999 |
| **Conditioning room males** | 0.003457138 | 6.0366E-06 | 0 | 2.76918E-05 |
| **Novel room males** | 0.835480603 | 0.000192999 | 2.76918E-05 | 0 |

##### 28 days

| | Conditioning room females | Novel room females | Conditioning room males | Novel room males |
|---|---|---|---|---|
| **Conditioning room females** | 1.1365E-233 | 0.06428691 | 1.96876E-19 | 0.000245406 |
| **Novel room females** | 0.06428691 | 0 | 0.039170721 | 5.83842E-05 |
| **Conditioning room males** | 1.96876E-19 | 0.039170721 | 0 | 1.00953E-05 |
| **Novel room males** | 0.000245406 | 5.83842E-05 | 1.00953E-05 | 0 |

##### 45 days

| | Conditioning room females | Novel room females | Conditioning room males | Novel room males |
|---|---|---|---|---|
| **Conditioning room females** | 0 | 0.00046589 | 0.040740727 | 0.19577853 |
| **Novel room females** | 0.00046589 | 0 | 0.00050415 | 0.000853511 |
| **Conditioning room males** | 0.040740727 | 0.00050415 | 0 | 0.539158623 |
| **Novel room males** | 0.19577853 | 0.000853511 | 0.539158623 | 0 |

#### Confidence Intervals

##### 2 days

| | Conditioning room females | Novel room females | Conditioning room males | Novel room males |
|---|---|---|---|---|
| **Conditioning room females** | 1.000 to 1.000 | 0.8114 to 0.9893 | 0.1886 to 0.7314 | -0.5523 to 0.6435 |
| **Novel room females** | 0.8114 to 0.9893 | 1.000 to 1.000 | 0.8561 to 0.9920 | 0.6779 to 0.9804 |
| **Conditioning room males** | 0.1886 to 0.7314 | 0.8561 to 0.9920 | 1.000 to 1.000 | 0.7566 to 0.9828 |
| **Novel room males** | -0.5523 to 0.6435 | 0.6779 to 0.9804 | 0.7566 to 0.9828 | 1.000 to 1.000 |

##### 28 days

| | Conditioning room females | Novel room females | Conditioning room males | Novel room males |
|---|---|---|---|---|
| **Conditioning room females** | 1.000 to 1.000 | -0.03581 to 0.8540 | 0.9249 to 0.9815 | 0.6214 to 0.9712 |
| **Novel room females** | -0.03581 to 0.8540 | 1.000 to 1.000 | 0.03978 to 0.8732 | 0.7163 to 0.9795 |
| **Conditioning room males** | 0.9249 to 0.9815 | 0.03978 to 0.8732 | 1.000 to 1.000 | 0.8027 to 0.9864 |
| **Novel room males** | 0.6214 to 0.9712 | 0.7163 to 0.9795 | 0.8027 to 0.9864 | 1.000 to 1.000 |

##### 45 days

| | Conditioning room females | Novel room females | Conditioning room males | Novel room males |
|---|---|---|---|---|
| **Conditioning room females** | 1.000 to 1.000 | 0.6529 to 0.9830 | 0.01911 to 0.6747 | -0.2379 to 0.8155 |
| **Novel room females** | 0.6529 to 0.9830 | 1.000 to 1.000 | 0.6460 to 0.9826 | 0.5964 to 0.9796 |
| **Conditioning room males** | 0.01911 to 0.6747 | 0.6460 to 0.9826 | 1.000 to 1.000 | -0.4476 to 0.7183 |
| **Novel room males** | -0.2379 to 0.8155 | 0.5964 to 0.9796 | -0.4476 to 0.7183 | 1.000 to 1.000 |

Correlation Matrices

**Pearson's r**

**2 days**

| | Curved female | Planar female | Curved male | Planar male |
|---|---|---|---|---|
| **Curved female** | 1.00 | 0.98 | 0.97 | 0.96 |
| **Planar female** | 0.98 | 1.00 | -0.01 | 0.58 |
| **Curved male** | 0.97 | -0.01 | 1.00 | 0.94 |
| **Planar male** | 0.96 | 0.58 | 0.94 | 1.00 |

**28 days**

| | Curved female | Planar female | Curved male | Planar male |
|---|---|---|---|---|
| **Curved female** | 1.00 | 0.94 | 0.97 | 0.98 |
| **Planar female** | 0.94 | 1.00 | 0.96 | 0.81 |
| **Curved male** | 0.97 | 0.96 | 1.00 | 0.94 |
| | 0.98 | 0.81 | 0.94 | 1.00 |

**45 days**

| | Curved female | Planar female | Curved male | Planar male |
|---|---|---|---|---|
| **Curved female** | 1.00 | 0.99 | 0.93 | 0.98 |
| **Planar female** | 0.99 | 1.00 | -0.15 | 0.63 |
| **Curved male** | 0.93 | -0.15 | 1.00 | -0.47 |
| **Planar male** | 0.98 | 0.63 | -0.47 | 1.00 |

**P-values**

**2 days**

| | Curved female | Planar female | Curved male | Planar male |
|---|---|---|---|---|
| **Curved female** | 0 | 4.23657E-07 | 3.87645E-06 | 8.21009E-06 |
| **Planar female** | 4.23657E-07 | 0 | 0.976512126 | 0.000565155 |
| **Curved male** | 3.87645E-06 | 0.976512126 | 0 | 1.23321E-05 |
| **Planar male** | 8.21009E-06 | 0.000565155 | 1.23321E-05 | 0 |

**28 days**

| | Curved female | Planar female | Curved male | Planar male |
|---|---|---|---|---|
| **Curved female** | 0 | 1.87238E-05 | 6.94195E-07 | 2.73913E-07 |
| **Planar female** | 1.87238E-05 | 1.0518E-244 | 1.89304E-06 | 9.50654E-09 |
| **Curved male** | 6.94195E-07 | 1.89304E-06 | 0 | 1.23321E-05 |
| **Planar male** | 2.73913E-07 | 9.50654E-09 | 1.23321E-05 | 0 |

**45 days**

| | Curved female | Planar female | Curved male | Planar male |
|---|---|---|---|---|
| **Curved female** | 0 | 4.32258E-07 | 0.000301501 | 2.63318E-06 |
| **Planar female** | 4.32258E-07 | 0 | 0.657896694 | 0.000374776 |
| **Curved male** | 0.000301501 | 0.657896694 | 0 | 0.145555803 |
| **Planar male** | 2.63318E-06 | 0.000374776 | 0.145555803 | 4.1612E-220 |

**Confidence Intervals**

**2 days**

| | Curved female | Planar female | Curved male | Planar male |
|---|---|---|---|---|
| **Curved female** | 1.000 to 1.000 | 0.9243 to 0.9959 | 0.8706 to 0.9929 | 0.8452 to 0.9914 |
| **Planar female** | 0.9243 to 0.9959 | 1.000 to 1.000 | -0.6063 to 0.5934 | 0.2893 to 0.7773 |
| **Curved male** | 0.8706 to 0.9929 | -0.6063 to 0.5934 | 1.000 to 1.000 | 0.7942 to 0.9858 |
| **Planar male** | 0.8452 to 0.9914 | 0.2893 to 0.7773 | 0.7942 to 0.9858 | 1.000 to 1.000 |

**28 days**

| | Curved female | Planar female | Curved male | Planar male |
|---|---|---|---|---|
| **Curved female** | 1.000 to 1.000 | 0.7755 to 0.9843 | 0.8882 to 0.9926 | 0.9086 to 0.9940 |
| **Planar female** | 0.7755 to 0.9843 | 1.000 to 1.000 | 0.8615 to 0.9907 | 0.6503 to 0.9036 |
| **Curved male** | 0.8882 to 0.9926 | 0.8615 to 0.9907 | 1.000 to 1.000 | 0.7942 to 0.9858 |
| **Planar male** | 0.9086 to 0.9940 | 0.6503 to 0.9036 | 0.7942 to 0.9858 | 1.000 to 1.000 |

**45 days**

| | Curved female | Planar female | Curved male | Planar male |
|---|---|---|---|---|
| **Curved female** | 1.000 to 1.000 | 0.9474 to 0.9978 | 0.6892 to 0.9851 | 0.9129 to 0.9963 |
| **Planar female** | 0.9474 to 0.9978 | 1.000 to 1.000 | -0.6884 to 0.4937 | 0.3359 to 0.8176 |
| **Curved male** | 0.6892 to 0.9851 | -0.6884 to 0.4937 | 1.000 to 1.000 | -0.8342 to 0.1820 |
| **Planar male** | 0.9129 to 0.9963 | 0.3359 to 0.8176 | -0.8342 to 0.1820 | 1.000 to 1.000 |

Correlation Matrices

| Pearson's r | | | | |
|---|---|---|---|---|
| **2 days** | | | | |
| | **Grid female** | **Smooth female** | **Grid male** | **Smooth male** |
| **Grid female** | 1.00 | 0.98 | 0.72 | 0.75 |
| **Smooth female** | 0.98 | 1.00 | 0.74 | 0.74 |
| **Grid male** | 0.72 | 0.74 | 1.00 | 0.96 |
| **Smooth male** | 0.75 | 0.74 | 0.96 | 1.00 |
| **28 days** | | | | |
| | **Grid female** | **Smooth female** | **Grid male** | **Smooth male** |
| **Grid female** | 1.00 | 0.94 | -0.29 | 0.95 |
| **Smooth female** | 0.94 | 1.00 | -0.47 | 0.98 |
| **Grid male** | -0.29 | -0.47 | 1.00 | -0.52 |
| **Smooth male** | 0.95 | 0.98 | -0.52 | 1.00 |
| **45 days** | | | | |
| | **Grid female** | **Smooth female** | **Grid male** | **Smooth male** |
| **Grid female** | 1.00 | 0.96 | 0.89 | 0.30 |
| **Smooth female** | 0.96 | 1.00 | 0.96 | 0.36 |
| **Grid male** | 0.89 | 0.96 | 1.00 | 0.35 |
| **Smooth male** | 0.30 | 0.36 | 0.35 | 1.00 |

| P-values | | | | |
|---|---|---|---|---|
| **2 days** | | | | |
| | **Grid female** | **Smooth female** | **Grid male** | **Smooth male** |
| **Grid female** | 0 | 4.50394E-14 | 0.000201288 | 9.0058E-05 |
| **Smooth female** | 4.50394E-14 | 0 | 0.000199732 | 0.000207776 |
| **Grid male** | 0.000201288 | 0.000199732 | 0 | 1.69531E-12 |
| **Smooth male** | 9.0058E-05 | 0.000207776 | 1.69531E-12 | 0 |
| **28 days** | | | | |
| | **Grid female** | **Smooth female** | **Grid male** | **Smooth male** |
| **Grid female** | 1.5156E-153 | 7.1847E-11 | 0.195225002 | 9.15513E-12 |
| **Smooth female** | 7.1847E-11 | 0 | 0.027631135 | 1.1363E-14 |
| **Grid male** | 0.195225002 | 0.027631135 | 0 | 0.013754196 |
| **Smooth male** | 9.15513E-12 | 1.1363E-14 | 0.013754196 | 1.5156E-153 |
| **45 days** | | | | |
| | **Grid female** | **Smooth female** | **Grid male** | **Smooth male** |
| **Grid female** | 0 | 4.16566E-10 | 7.07757E-07 | 0.225130444 |
| **Smooth female** | 4.16566E-10 | 2.2665E-121 | 1.24325E-10 | 0.141679385 |
| **Grid male** | 7.07757E-07 | 1.24325E-10 | 0 | 0.128855033 |
| **Smooth male** | 0.225130444 | 0.141679385 | 0.128855033 | 0 |

| Confidence Intervals of r | | | | |
|---|---|---|---|---|
| **2 days** | | | | |
| | **Grid female** | **Smooth female** | **Grid male** | **Smooth male** |
| **Grid female** | 1.0 to 1.0 | 0.95 to 0.99 | 0.43 to 0.88 | 0.47 to 0.89 |
| **Smooth female** | 0.95 to 0.99 | 1.0 to 1.0 | 0.44 to 0.89 | 0.44 to 0.89 |
| **Grid male** | 0.43 to 0.88 | 0.44 to 0.89 | 1.0 to 1.0 | 0.90 to 0.98 |
| **Smooth male** | 0.47 to 0.89 | 0.44 to 0.89 | 0.90 to 0.98 | 1.0 to 1.0 |
| **28 days** | | | | |
| | **Grid female** | **Smooth female** | **Grid male** | **Smooth male** |
| **Grid female** | 1.000 to 1.000 | 0.8610 to 0.9756 | -0.6322 to 0.1531 | 0.8866 to 0.9803 |
| **Smooth female** | 0.8610 to 0.9756 | 1.000 to 1.000 | -0.7436 to -0.05922 | 0.9415 to 0.9901 |
| **Grid male** | -0.6322 to 0.1531 | -0.7436 to -0.05922 | 1.000 to 1.000 | -0.7706 to -0.1219 |
| **Smooth male** | 0.8866 to 0.9803 | 0.9415 to 0.9901 | -0.7706 to -0.1219 | 1.000 to 1.000 |
| **45 days** | | | | |
| | **Grid female** | **Smooth female** | **Grid male** | **Smooth male** |
| **Grid female** | 1.000 to 1.000 | 0.8889 to 0.9846 | 0.7262 to 0.9590 | -0.1932 to 0.6732 |
| **Smooth female** | 0.8889 to 0.9846 | 1.000 to 1.000 | 0.9042 to 0.9868 | -0.1279 to 0.7082 |
| **Grid male** | 0.7262 to 0.9590 | 0.9042 to 0.9868 | 1.000 to 1.000 | -0.1081 to 0.6870 |
| **Smooth male** | -0.1932 to 0.6732 | -0.1279 to 0.7082 | -0.1081 to 0.6870 | 1.000 to 1.000 |

Correlation Matrices

| Pearson's r | | | | | | |
|---|---|---|---|---|---|---|
| **2 days** | | | | | | |
| | **Ethanol female** | **Isopropyl female** | **QA female** | **Ethanol male** | **Isopropyl male** | **QA male** |
| **Ethanol female** | 1.00 | 0.20 | 0.95 | 0.62 | -0.19 | 0.07 |
| **Isopropyl female** | 0.20 | 1.00 | 0.98 | 0.96 | 0.90 | 0.96 |
| **QA female** | 0.95 | 0.98 | 1.00 | 0.97 | 0.99 | 0.92 |
| **Ethanol male** | 0.62 | 0.96 | 0.97 | 1.00 | 0.90 | 0.93 |
| **Isopropyl male** | -0.19 | 0.90 | 0.99 | 0.90 | 1.00 | 0.92 |
| **QA male** | 0.07 | 0.96 | 0.92 | 0.93 | 0.92 | 1.00 |
| **28 days** | | | | | | |
| | **Ethanol female** | **Isopropyl female** | **QA female** | **Ethanol male** | **Isopropyl male** | **QA male** |
| **Ethanol female** | 1.00 | 0.91 | 0.55 | 0.95 | 0.92 | 0.89 |
| **Isopropyl female** | 0.91 | 1.00 | 0.95 | 0.95 | 0.98 | 0.99 |
| **QA female** | 0.55 | 0.95 | 1.00 | 0.60 | 0.93 | 0.92 |
| **Ethanol male** | 0.95 | 0.95 | 0.60 | 1.00 | 0.97 | 0.95 |
| **Isopropyl male** | 0.92 | 0.98 | 0.93 | 0.97 | 1.00 | 0.96 |
| **QA male** | 0.89 | 0.99 | 0.92 | 0.95 | 0.96 | 1.00 |
| **45 days** | | | | | | |
| | **Ethanol female** | **Isopropyl female** | **QA female** | **Ethanol male** | **Isopropyl male** | **QA male** |
| **Ethanol female** | 1.00 | 0.92 | 0.92 | 0.87 | 0.10 | 0.42 |
| **Isopropyl female** | 0.92 | 1.00 | 0.97 | 0.94 | 0.83 | 0.91 |
| **QA female** | 0.92 | 0.97 | 1.00 | 0.92 | 0.83 | 0.90 |
| **Ethanol male** | 0.87 | 0.94 | 0.92 | 1.00 | -0.07 | 0.21 |
| **Isopropyl male** | 0.10 | 0.83 | 0.83 | -0.07 | 1.00 | 0.89 |
| **QA male** | 0.42 | 0.91 | 0.90 | 0.21 | 0.89 | 1.00 |

| P-values | | | | | | |
|---|---|---|---|---|---|---|
| **2 days** | | | | | | |
| | **Ethanol female** | **Isopropyl female** | **QA female** | **Ethanol male** | **Isopropyl male** | **QA male** |
| **Ethanol female** | 0 | 0.563206614 | 1.888E-05 | 0.003384902 | 0.579874559 | 0.8354806 |
| **Isopropyl female** | 0.563206614 | 0 | 8.259E-07 | 1.62308E-06 | 0.000148501 | 3.3779E-06 |
| **QA female** | 1.88803E-05 | 8.25903E-07 | 0 | 6.0366E-06 | 8.92579E-08 | 0.000193 |
| **Ethanol male** | 0.003384902 | 1.62308E-06 | 6.0366E-06 | 0 | 0.000180086 | 2.7692E-05 |
| **Isopropyl male** | 0.579874559 | 0.000148501 | 8.9258E-08 | 0.000180086 | 0 | 5.4895E-05 |
| **QA male** | 0.835480603 | 3.37791E-06 | 0.000193 | 2.76918E-05 | 5.48952E-05 | 0 |
| **28 days** | | | | | | |
| | **Ethanol female** | **Isopropyl female** | **QA female** | **Ethanol male** | **Isopropyl male** | **QA male** |
| **Ethanol female** | 0 | 0.000127028 | 0.06428691 | 1.97506E-11 | 5.84085E-05 | 0.00024541 |
| **Isopropyl female** | 0.000127028 | 0 | 7.151E-06 | 6.47485E-06 | 3.00337E-07 | 3.507E-08 |
| **QA female** | 0.06428691 | 7.151E-06 | 0 | 0.039170721 | 3.74194E-05 | 5.8384E-05 |
| **Ethanol male** | 1.97506E-11 | 6.47485E-06 | 0.03917072 | 0 | 5.63649E-07 | 1.0095E-05 |
| **Isopropyl male** | 5.84085E-05 | 3.00337E-07 | 3.7419E-05 | 5.63649E-07 | 0 | 2.9081E-06 |
| **QA male** | 0.000245406 | 3.50697E-08 | 5.8384E-05 | 1.00953E-05 | 2.90808E-06 | 0 |
| **45 days** | | | | | | |
| | **Ethanol female** | **Isopropyl female** | **QA female** | **Ethanol male** | **Isopropyl male** | **QA male** |
| **Ethanol female** | 0 | 0.000424545 | 0.00046589 | 2.99982E-06 | 0.771367907 | 0.19577853 |
| **Isopropyl female** | 0.000424545 | 0 | 1.282E-05 | 0.000194989 | 0.005194189 | 0.00066432 |
| **QA female** | 0.00046589 | 1.28202E-05 | 0 | 0.00050415 | 0.005196754 | 0.00085351 |
| **Ethanol male** | 2.99982E-06 | 0.000194989 | 0.00050415 | 0 | 0.844925787 | 0.53915862 |
| **Isopropyl male** | 0.771367907 | 0.005194189 | 0.00519675 | 0.844925787 | 0 | 0.00023498 |
| **QA male** | 0.19577853 | 0.000664325 | 0.00085351 | 0.539158623 | 0.000234976 | 0 |

Correlation Matrices

| Confidence Intervals of r | | | | | | |
|---|---|---|---|---|---|---|
| **2 days** | | | | | | |
| | **Ethanol female** | **Isopropyl female** | **QA female** | **Ethanol male** | **Isopropyl male** | **QA male** |
| **Ethanol female** | 1.000 to 1.000 | -0.4576 to 0.7122 | 0.8114 to 0.9893 | 0.2482 to 0.8349 | -0.7080 to 0.4642 | -0.5523 to 0.6435 |
| **Isopropyl female** | -0.4576 to 0.7122 | 1.000 to 1.000 | 0.9109 to 0.9952 | 0.8659 to 0.9911 | 0.6574 to 0.9745 | 0.8433 to 0.9894 |
| **QA female** | 0.8114 to 0.9893 | 0.9109 to 0.9952 | 1.000 to 1.000 | 0.8561 to 0.9920 | 0.9483 to 0.9973 | 0.6779 to 0.9804 |
| **Ethanol male** | 0.2482 to 0.8349 | 0.8659 to 0.9911 | 0.8561 to 0.9920 | 1.000 to 1.000 | 0.6440 to 0.9733 | 0.7566 to 0.9828 |
| **Isopropyl male** | -0.7080 to 0.4642 | 0.6574 to 0.9745 | 0.9483 to 0.9973 | 0.6440 to 0.9733 | 1.000 to 1.000 | 0.7199 to 0.9798 |
| **QA male** | -0.5523 to 0.6435 | 0.8433 to 0.9894 | 0.6779 to 0.9804 | 0.7566 to 0.9828 | 0.7199 to 0.9798 | 1.000 to 1.000 |
| **28 days** | | | | | | |
| | **Ethanol female** | **Isopropyl female** | **QA female** | **Ethanol male** | **Isopropyl male** | **QA male** |
| **Ethanol female** | 1.000 to 1.000 | 0.6680 to 0.9754 | -0.03581 to 0.8540 | 0.8776 to 0.9787 | 0.7163 to 0.9795 | 0.6214 to 0.9712 |
| **Isopropyl female** | 0.6680 to 0.9754 | 1.000 to 1.000 | 0.8164 to 0.9874 | 0.8202 to 0.9877 | 0.9067 to 0.9939 | 0.9415 to 0.9962 |
| **QA female** | -0.03581 to 0.8540 | 0.8164 to 0.9874 | 1.000 to 1.000 | 0.03978 to 0.8732 | 0.7410 to 0.9816 | 0.7163 to 0.9795 |
| **Ethanol male** | 0.8776 to 0.9787 | 0.8202 to 0.9877 | 0.03978 to 0.8732 | 1.000 to 1.000 | 0.8931 to 0.9930 | 0.8027 to 0.9864 |
| **Isopropyl male** | 0.7163 to 0.9795 | 0.9067 to 0.9939 | 0.7410 to 0.9816 | 0.8931 to 0.9930 | 1.000 to 1.000 | 0.8482 to 0.9898 |
| **QA male** | 0.6214 to 0.9712 | 0.9415 to 0.9962 | 0.7163 to 0.9795 | 0.8027 to 0.9864 | 0.8482 to 0.9898 | 1.000 to 1.000 |
| **45 days** | | | | | | |
| | **Ethanol female** | **Isopropyl female** | **QA female** | **Ethanol male** | **Isopropyl male** | **QA male** |
| **Ethanol female** | 1.000 to 1.000 | 0.6610 to 0.9835 | 0.6529 to 0.9830 | 0.6749 to 0.9500 | -0.5323 to 0.6599 | -0.2379 to 0.8155 |
| **Isopropyl female** | 0.6610 to 0.9835 | 1.000 to 1.000 | 0.8654 to 0.9941 | 0.7222 to 0.9869 | 0.3808 to 0.9641 | 0.6207 to 0.9811 |
| **QA female** | 0.6529 to 0.9830 | 0.8654 to 0.9941 | 1.000 to 1.000 | 0.6460 to 0.9826 | 0.3808 to 0.9641 | 0.5964 to 0.9796 |
| **Ethanol male** | 0.6749 to 0.9500 | 0.7222 to 0.9869 | 0.6460 to 0.9826 | 1.000 to 1.000 | -0.6411 to 0.5552 | -0.4476 to 0.7183 |
| **Isopropyl male** | -0.5323 to 0.6599 | 0.3808 to 0.9641 | 0.3808 to 0.9641 | -0.6411 to 0.5552 | 1.000 to 1.000 | 0.6246 to 0.9715 |
| **QA male** | -0.2379 to 0.8155 | 0.6207 to 0.9811 | 0.5964 to 0.9796 | -0.4476 to 0.7183 | 0.6246 to 0.9715 | 1.000 to 1.000 |

**Supplemental Table 2.** Pearson's correlation coefficients and their respective p-values and confidence intervals for each interval (parameter variables vs. sex). "QA": quaternary ammonium.

#### *PCAs - Multivariate Analysis of context parameters and some of their variants*

In order to reduce dimensionality of our data with minimal information loss and thus increase the interpretability of our results, we performed Principal Component Analysis (PCA) of our heatmap datasets. **Figure S-4** below shows the results for PCAs and Scree Plots: (A, B) for the four contexts A, B, C and D in the main experiment organized per sex and training-test interval time point, (C, D) for the 11 variables of the 5 selected parameters compared simultaneously, and (E.F) for the 5 parameters analyzed separately in terms of their variables and sex.

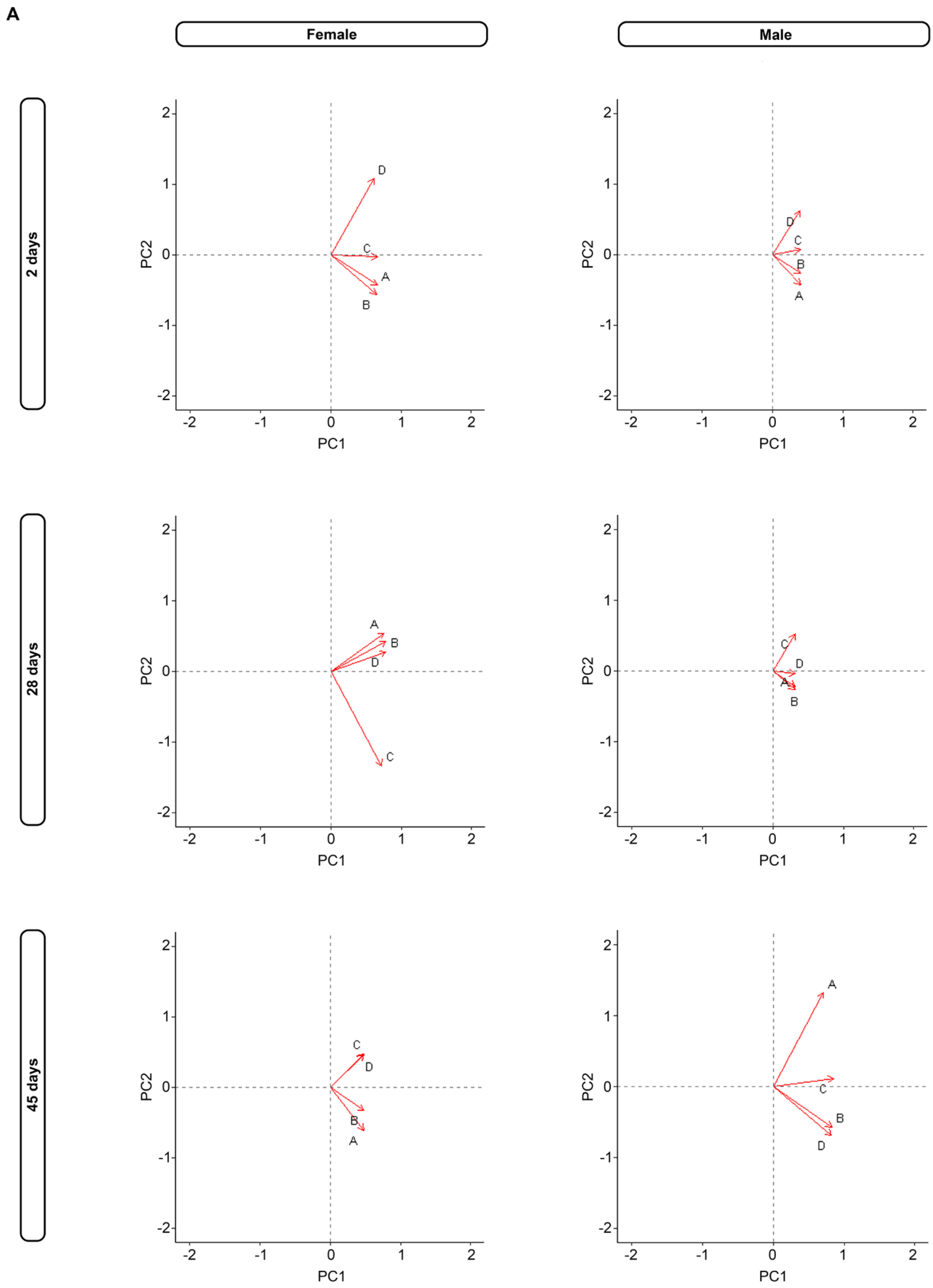

A
Female
Male
2 days
28 days
45 days
PC1
PC2
A
B
C
D

B

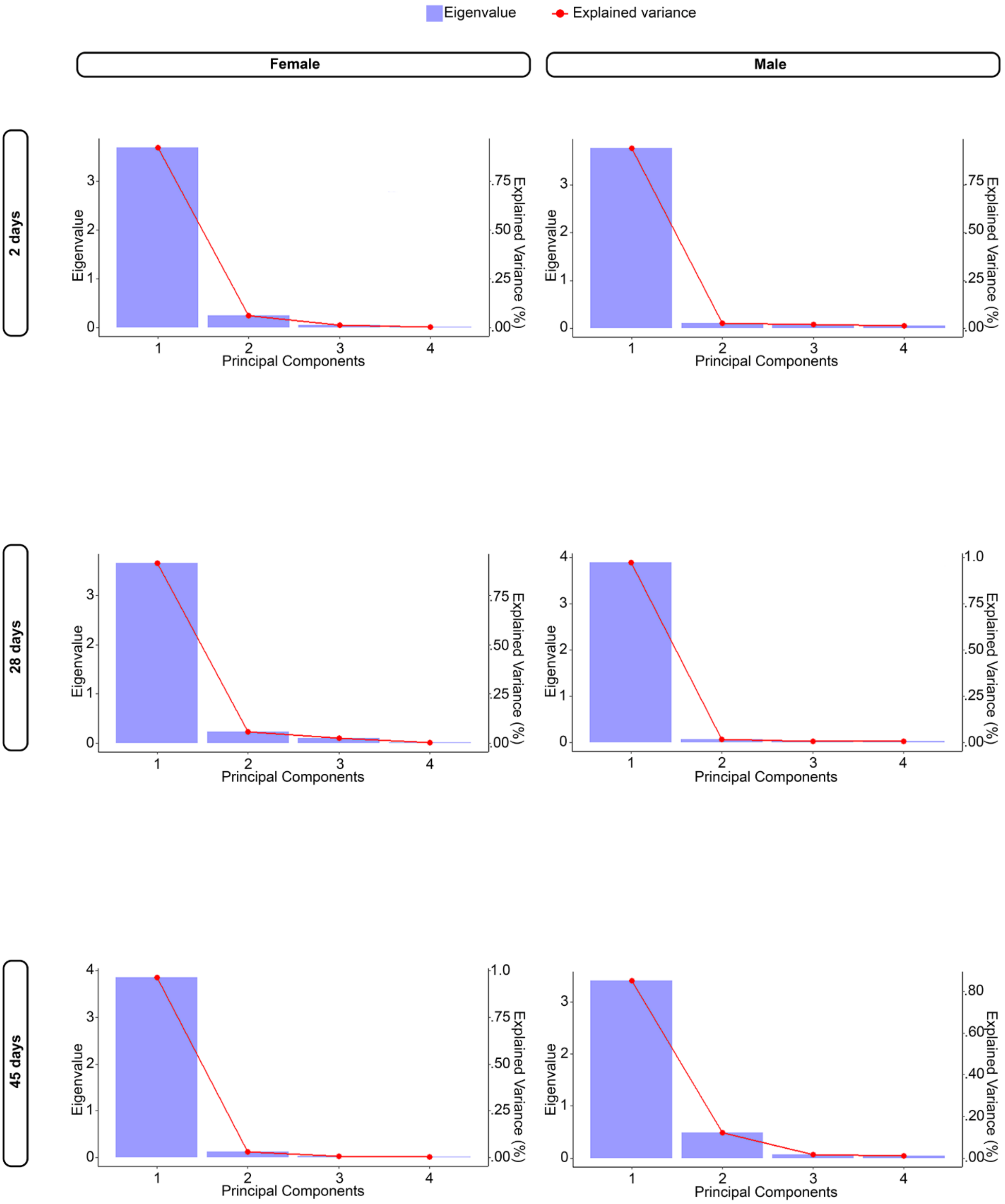

Eigenvalue
Explained variance
Female
Male
2 days
28 days
45 days
Eigenvalue
Explained Variance (%)
Principal Components

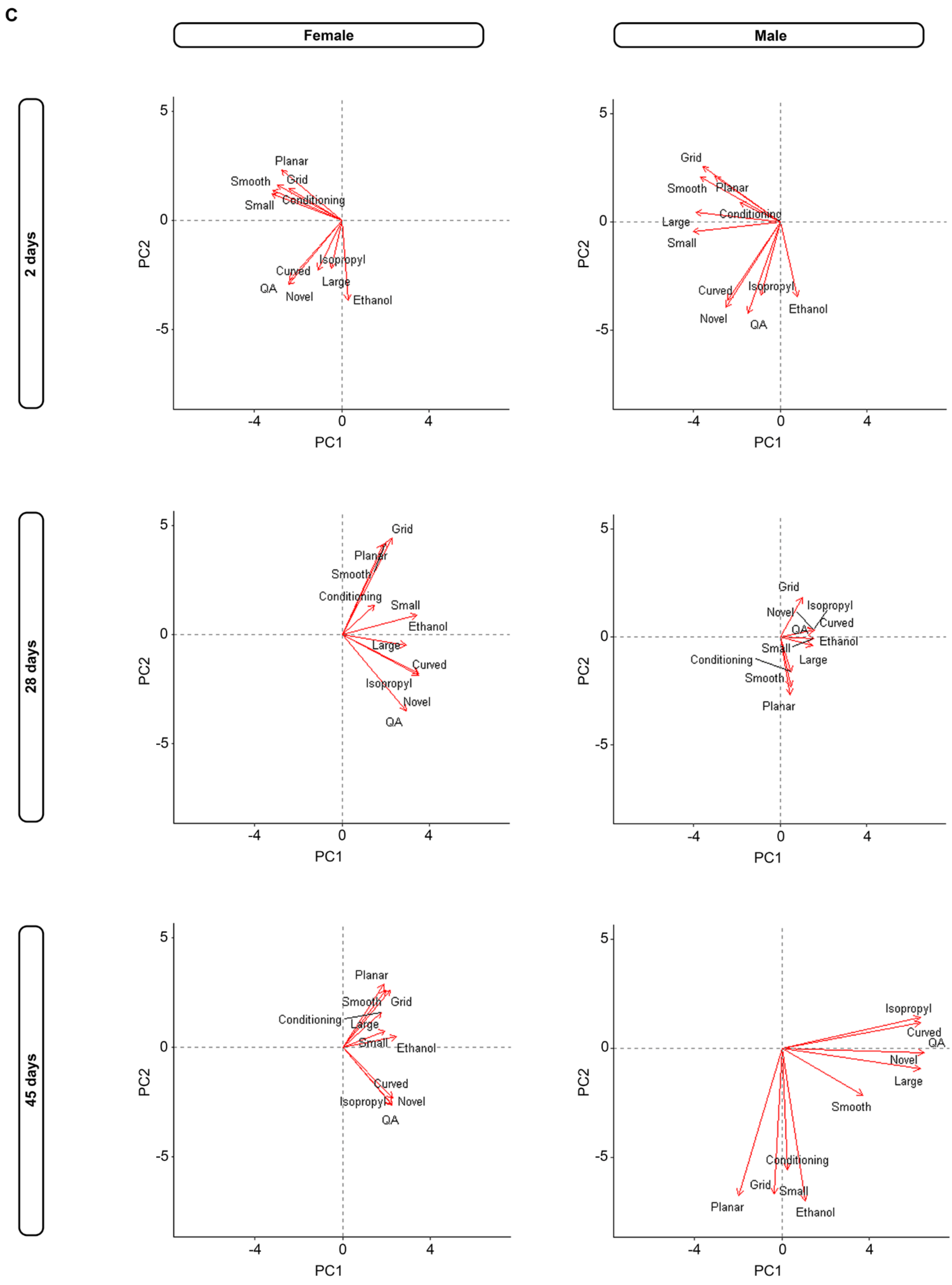
C
Female
Male
2 days
28 days
45 days
PC1
PC2
Planar
Smooth
Grid
Small
Conditioning
Isopropyl
Curved
QA
Large
Novel
Ethanol
Large

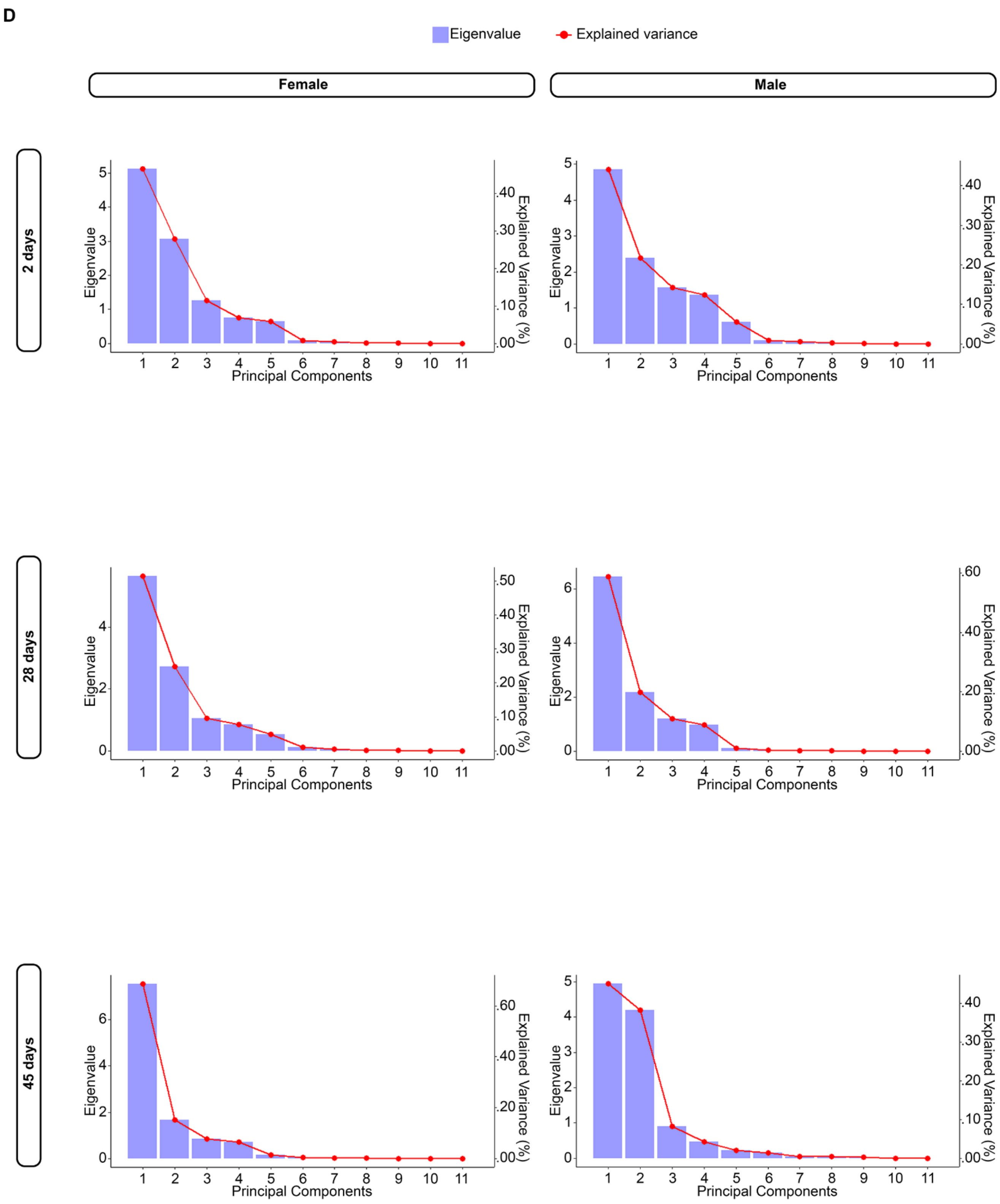
D
Eigenvalue
Explained variance
Female
Male
2 days
28 days
45 days
Eigenvalue
Explained Variance (%)
Principal Components

E

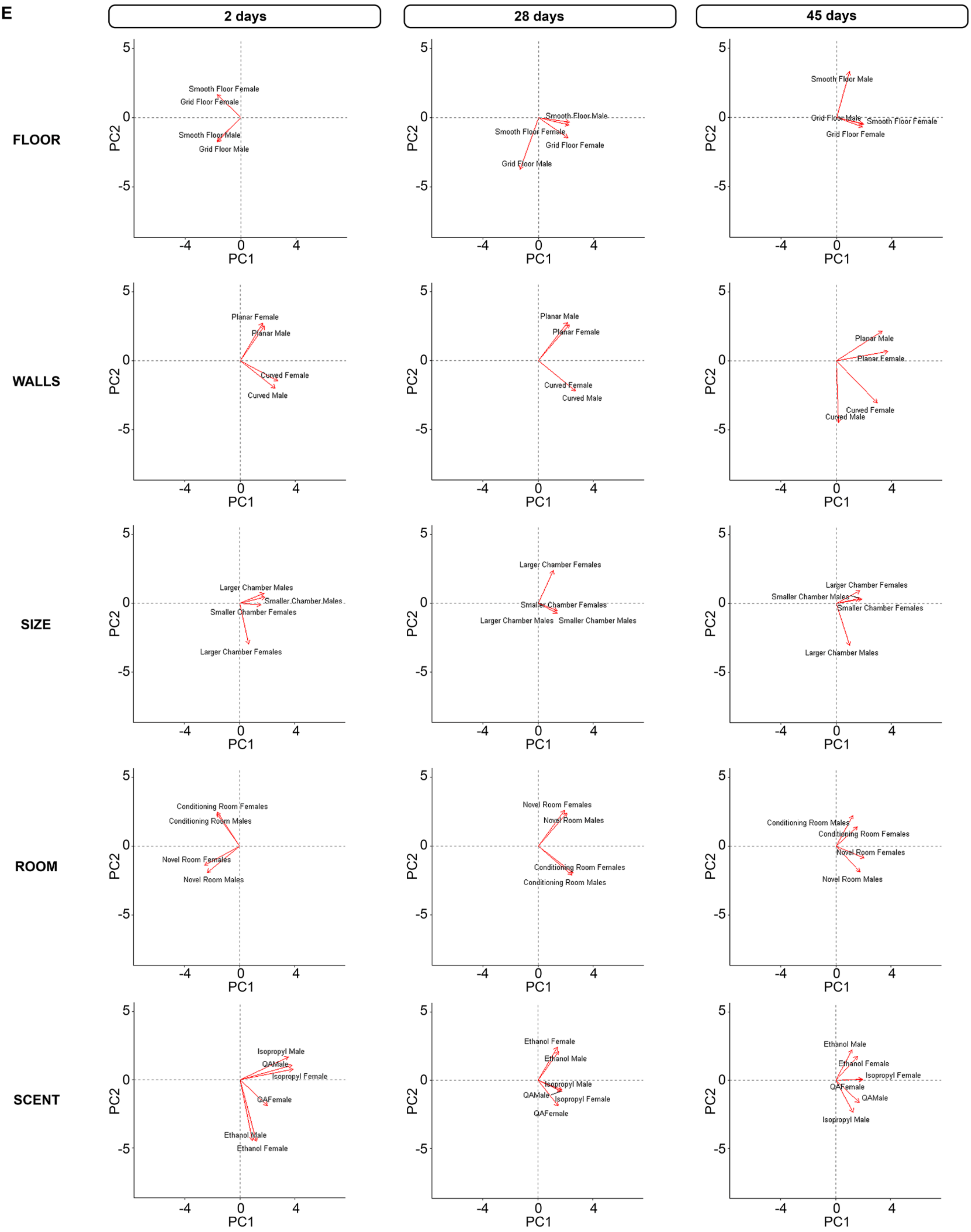
2 days
28 days
45 days
FLOOR
WALLS
SIZE
ROOM
SCENT
PC1
PC2
Smooth Floor Female
Grid Floor Female
Smooth Floor Male
Grid Floor Male
Smooth Floor Male
Smooth Floor Female
Grid Floor Female
Grid Floor Male
Smooth Floor Male
Grid Floor Male
Smooth Floor Female
Grid Floor Female
Planar Female
Planar Male
Curved Female
Curved Male
Planar Male
Planar Female
Curved Female
Curved Male
Planar Male
Planar Female
Curved Female
Curved Male
Larger Chamber Males
Smaller Chamber Males
Smaller Chamber Females
Larger Chamber Females
Larger Chamber Females
Smaller Chamber Females
Larger Chamber Males
Smaller Chamber Males
Larger Chamber Females
Smaller Chamber Males
Smaller Chamber Females
Larger Chamber Males
Conditioning Room Females
Conditioning Room Males
Novel Room Females
Novel Room Males
Novel Room Females
Novel Room Males
Conditioning Room Females
Conditioning Room Males
Conditioning Room Males
Conditioning Room Females
Novel Room Females
Novel Room Males
Isopropyl Male
QAMale
Isopropyl Female
QAFemale
Ethanol Male
Ethanol Female
Ethanol Female
Ethanol Male
Isopropyl Male
QAMale
Isopropyl Female
QAFemale
Ethanol Male
Ethanol Female
Isopropyl Female
QAFemale
QAMale
Isopropyl Male

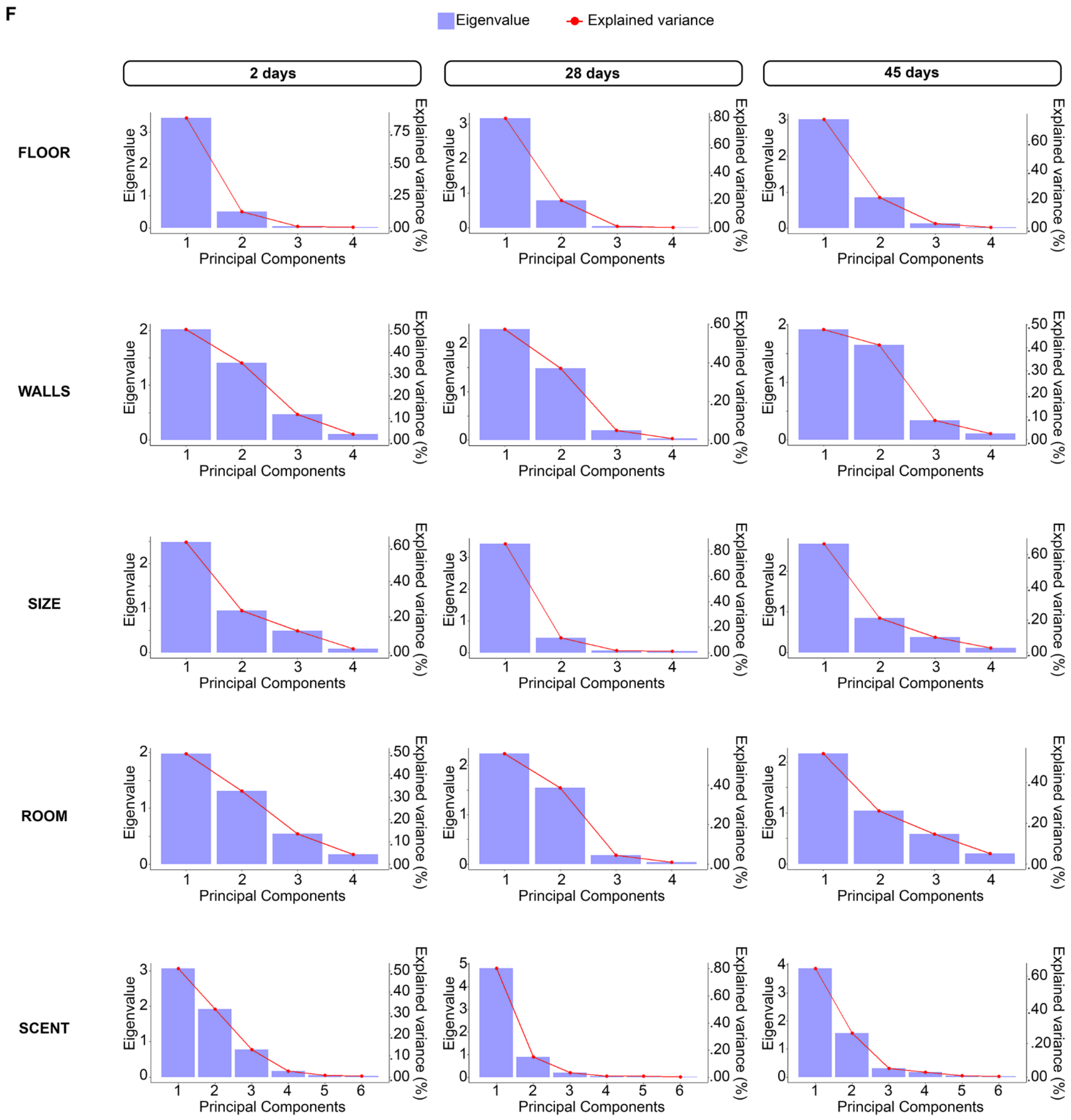


**Supplemental Figure 4.** Bidimensional Principal Component Analysis (PCA) loading and scree plots comparing different Main Experiment variables for different sexes and training-test intervals. **A:** Loading plots for the four contexts organized by sex and training-test interval. **B:** Scree plots showing explained variance (lines) and eigenvalues (bars) for the four contexts. **C:** Loading plots with parameter variables (grid vs. smooth, curved vs. planar, larger vs. smaller, conditioning vs. novel, ethanol vs. isopropyl alcohol vs. quaternary ammonium) compared across sexes and different intervals. **D:** Scree plots showing explained variance (lines) and eigenvalues (bars) for the different variable groupings. **E:** Loading plots grouped by sex, comparing the different sensory modality parameters (Cleaning agent scent, Wall geometry, Test session room, Floor texture and

Test chamber size) for each interval. **F:** Scree plots showing explained variance (lines) and eigenvalues (bars) for the different parameters grouped by sex. In all PCA plots the length of eigenvectors (arrows) indicates the magnitude of the loading for each modality. Inclination < 90° (positive loadings) indicates correlated variables, > 90° (negative loadings) indicates inversely correlated variables and = 90°, independent variables. Pooled contexts for each sensory modality are the same as in the heatmaps: Smooth floor (contexts B and C), Grid floor (contexts A and D), Curved walls (context B) and Planar walls (contexts A, C and D), Larger test chamber (contexts C and D) and Smaller test chamber (contexts A and B), Conditioning room (contexts A, B and D) and Novel room (context C), 70% ethanol (contexts A and B), 70% isopropyl (context D) and quaternary ammonium (context C). QA: quaternary ammonium.

### PCA loading scores

| 2 days | | | | | | | | | | | | | | | | | | | | | |
|---|---|---|---|---|---|---|---|---|---|---|---|---|---|---|---|---|---|---|---|---|---|
| Chamber Size | | | | Wall Geometry | | | | Cleaning Agent Scent | | | | | | Floor Texture | | | | Test Session Room | | | |
| **PC1** | **PC2** | **PC3** | **PC4** | **PC1** | **PC2** | **PC3** | **PC4** | **PC1** | **PC2** | **PC3** | **PC4** | **PC5** | **PC6** | **PC1** | **PC2** | **PC3** | **PC4** | **PC1** | **PC2** | **PC3** | **PC4** |
| -1.018 | 0.792 | 0.364 | -0.278 | -3.124 | 3.032 | -0.349 | 0.129 | 3.026 | 4.151 | -1.968 | -0.100 | 0.306 | 0.126 | -0.303 | -0.387 | 0.079 | -0.355 | 2.522 | 2.217 | 0.142 | -0.065 |
| -0.703 | 0.797 | 0.197 | -0.446 | -2.616 | 2.749 | -0.343 | 0.128 | -3.714 | 1.610 | -0.073 | -0.009 | -0.040 | -0.285 | -0.440 | -0.366 | 0.086 | -0.260 | 1.760 | 1.712 | 0.285 | -0.434 |
| -0.446 | 0.769 | 0.516 | -0.411 | -0.990 | 1.933 | -0.157 | 0.284 | -3.000 | 1.189 | -0.351 | 0.106 | -0.371 | 0.078 | -0.926 | -0.173 | -0.016 | -0.078 | 1.577 | 1.879 | 0.088 | -0.464 |
| -0.034 | 0.710 | 0.722 | -0.220 | -0.166 | 1.701 | -0.007 | 0.249 | -2.812 | 0.870 | -0.207 | 0.118 | -0.302 | 0.080 | -1.254 | -0.036 | -0.030 | -0.083 | 1.187 | 1.972 | -0.102 | -0.329 |
| 0.662 | 0.352 | 0.503 | 0.265 | 1.011 | 1.296 | 0.160 | -0.357 | -2.131 | 0.312 | 0.876 | -0.068 | 0.177 | 0.356 | -1.676 | 0.085 | 0.167 | 0.099 | -0.137 | 1.524 | -0.417 | 0.734 |
| 1.000 | 0.286 | 0.858 | 0.256 | 1.720 | 1.133 | 0.403 | -0.369 | -1.174 | 0.337 | 0.811 | -0.120 | 0.468 | -0.018 | -1.980 | 0.282 | 0.098 | 0.049 | -0.527 | 1.605 | -0.662 | 0.712 |
| 1.874 | 0.573 | 0.353 | 0.238 | 2.188 | 1.019 | 0.268 | -0.427 | -0.656 | 0.383 | 0.721 | -0.162 | 0.452 | -0.374 | -2.163 | 0.208 | 0.215 | 0.079 | -1.744 | 0.823 | -0.034 | -0.543 |
| 2.080 | 0.254 | 0.336 | 0.235 | 2.392 | 0.955 | 0.267 | -0.331 | 0.973 | 0.482 | 1.309 | 0.006 | -0.061 | 0.445 | -2.242 | 0.223 | 0.192 | 0.063 | -2.614 | 0.418 | -0.352 | 0.166 |
| 2.427 | 0.130 | 0.348 | 0.032 | 3.116 | 0.573 | 0.242 | 0.034 | 1.942 | 0.378 | 1.714 | 0.084 | -0.071 | -0.251 | -2.451 | 0.255 | 0.089 | 0.102 | -3.555 | -0.073 | -0.436 | 0.078 |
| 2.668 | 0.050 | 0.498 | 0.075 | 3.646 | 0.368 | 0.220 | 0.375 | 2.721 | 0.412 | 1.755 | 0.167 | -0.159 | 0.032 | -2.638 | 0.285 | -0.002 | 0.066 | -3.911 | -0.095 | -0.599 | 0.199 |
| 0.929 | 1.812 | -2.522 | 0.325 | 1.720 | -1.096 | -2.455 | 1.078 | 2.220 | 1.008 | -0.255 | 0.016 | -0.551 | -0.243 | 0.353 | -2.899 | -0.089 | 0.104 | -1.758 | -1.176 | 2.085 | -1.375 |
| -2.987 | 0.425 | -0.144 | 0.023 | -0.751 | -1.177 | -0.169 | -0.195 | -0.071 | 0.312 | 0.128 | -0.061 | -0.010 | -0.001 | 3.201 | 0.135 | 0.244 | -0.031 | 0.277 | -0.430 | 0.570 | 0.268 |
| -2.628 | 0.411 | 0.085 | 0.040 | -0.644 | -1.024 | -0.333 | -0.243 | 0.038 | -0.149 | -0.042 | -0.139 | -0.010 | 0.002 | 2.929 | -0.072 | 0.016 | 0.080 | 0.147 | -0.231 | 0.425 | 0.189 |
| -2.205 | 0.459 | 0.093 | 0.387 | -0.621 | -0.984 | -0.293 | -0.223 | 0.134 | -0.545 | -0.184 | -0.253 | -0.014 | 0.005 | 2.859 | -0.014 | 0.054 | 0.137 | 0.024 | -0.047 | 0.443 | 0.173 |
| -1.610 | -0.130 | 0.184 | 0.324 | -0.497 | -0.778 | -0.101 | -0.125 | 0.191 | -0.793 | -0.286 | -0.202 | -0.006 | 0.005 | 2.287 | 0.406 | 0.172 | 0.182 | -0.118 | 0.171 | 0.295 | 0.090 |
| -1.066 | -0.497 | 0.039 | 0.107 | -0.459 | -0.719 | -0.106 | -0.120 | 0.272 | -1.104 | -0.368 | -0.548 | -0.031 | 0.010 | 1.914 | 0.433 | 0.209 | -0.022 | -0.190 | 0.279 | 0.246 | 0.059 |
| -0.766 | -0.759 | -0.032 | 0.099 | -0.444 | -0.698 | -0.132 | -0.128 | 0.282 | -1.154 | -0.394 | -0.494 | -0.026 | 0.010 | 1.736 | 0.461 | 0.160 | -0.161 | -0.239 | 0.353 | 0.233 | 0.045 |
| -0.297 | -1.593 | -0.324 | -0.076 | -0.144 | -0.251 | -0.367 | -0.172 | 0.393 | -1.671 | -0.641 | -0.088 | 0.016 | 0.009 | 0.693 | 0.426 | -0.361 | -0.071 | -0.261 | 0.386 | 0.247 | 0.046 |
| -0.082 | -1.749 | -0.343 | -0.121 | -0.081 | -0.150 | -0.312 | -0.140 | 0.438 | -1.852 | -0.701 | -0.180 | 0.010 | 0.010 | 0.448 | 0.532 | -0.320 | -0.014 | -0.300 | 0.446 | 0.217 | 0.027 |
| 0.283 | -1.670 | -0.493 | -0.071 | 0.028 | 0.012 | -0.420 | -0.165 | 0.486 | -2.040 | -0.757 | -0.335 | 0.000 | 0.012 | 0.130 | 0.346 | -0.422 | 0.113 | -0.354 | 0.525 | 0.262 | 0.035 |
| 0.646 | -1.888 | -0.458 | 0.013 | -0.330 | -0.591 | -1.061 | -0.484 | 0.200 | -0.967 | -0.492 | 1.024 | 0.101 | -0.004 | 0.016 | 0.364 | -0.483 | -0.028 | 0.461 | -0.731 | 1.672 | 0.721 |
| 1.275 | 0.467 | -0.781 | -0.797 | -0.205 | -0.408 | -1.195 | -0.518 | 0.242 | -1.168 | -0.594 | 1.237 | 0.122 | -0.005 | -0.493 | -0.495 | -0.056 | 0.029 | 0.402 | -0.643 | 1.677 | 0.712 |
| | | | | -1.111 | -1.691 | 0.413 | -0.020 | | | | | | | | | | | 1.459 | -2.184 | -0.248 | 0.173 |
| | | | | -1.074 | -1.638 | 0.371 | -0.032 | | | | | | | | | | | 1.422 | -2.130 | -0.208 | 0.182 |
| | | | | -0.909 | -1.362 | 0.633 | 0.102 | | | | | | | | | | | 1.218 | -1.816 | -0.521 | 0.025 |
| | | | | -0.730 | -1.077 | 0.730 | 0.172 | | | | | | | | | | | 1.011 | -1.502 | -0.656 | -0.064 |
| | | | | -0.532 | -0.776 | 0.651 | 0.173 | | | | | | | | | | | 0.798 | -1.184 | -0.604 | -0.083 |
| | | | | -0.266 | -0.340 | 0.963 | 0.344 | | | | | | | | | | | 0.478 | -0.694 | -0.988 | -0.288 |
| | | | | -0.133 | -0.138 | 0.894 | 0.339 | | | | | | | | | | | 0.336 | -0.482 | -0.936 | -0.295 |
| | | | | -0.091 | -0.078 | 0.844 | 0.325 | | | | | | | | | | | 0.294 | -0.420 | -0.887 | -0.284 |
| | | | | 0.044 | 0.131 | 0.832 | 0.344 | | | | | | | | | | | 0.144 | -0.196 | -0.897 | -0.315 |
| | | | | -0.196 | -0.278 | 0.339 | 0.104 | | | | | | | | | | | 0.310 | -0.452 | -0.549 | -0.152 |
| | | | | 0.248 | 0.351 | -0.429 | -0.131 | | | | | | | | | | | -0.119 | 0.174 | 0.211 | 0.059 |

| 28 days | | | | | | | | | | | | | | | | | | | | | |
|---|---|---|---|---|---|---|---|---|---|---|---|---|---|---|---|---|---|---|---|---|---|
| Chamber Size | | | | Wall Geometry | | | | Cleaning Agent Scent | | | | | | Floor Texture | | | | Test Session Room | | | |
| **PC1** | **PC2** | **PC3** | **PC4** | **PC1** | **PC2** | **PC3** | **PC4** | **PC1** | **PC2** | **PC3** | **PC4** | **PC5** | **PC6** | **PC1** | **PC2** | **PC3** | **PC4** | **PC1** | **PC2** | **PC3** | **PC4** |
| -3.485 | 0.306 | -0.584 | 0.020 | -2.964 | 3.099 | 0.301 | 0.011 | -5.127 | -0.288 | 0.033 | 0.321 | -0.308 | -0.192 | 0.870 | 2.021 | 0.084 | 0.009 | -2.392 | -2.429 | -0.364 | -0.265 |
| -2.594 | 0.084 | 0.203 | -0.557 | -1.987 | 2.817 | -0.003 | -0.335 | -4.025 | 0.201 | 0.104 | -0.515 | -0.103 | -0.090 | 1.204 | 1.545 | 0.207 | -0.204 | -1.774 | -2.318 | -0.242 | 0.103 |
| -1.657 | -0.245 | 0.358 | 0.101 | -0.920 | 2.126 | -0.085 | 0.335 | -2.963 | 0.765 | 0.133 | 0.037 | 0.163 | 0.111 | 1.402 | 1.133 | 0.227 | 0.001 | -1.253 | -2.246 | -0.187 | -0.022 |
| -1.167 | -0.333 | 0.285 | -0.102 | -0.140 | 1.700 | -0.068 | 0.200 | -2.135 | 0.649 | -0.075 | -0.123 | 0.080 | 0.247 | 1.494 | 0.869 | 0.146 | -0.051 | -0.717 | -1.786 | 0.155 | 0.005 |
| -0.710 | -0.246 | 0.081 | -0.082 | 0.412 | 1.514 | 0.108 | 0.252 | -0.697 | 0.095 | -0.464 | -0.099 | 0.425 | -0.118 | 1.430 | 0.338 | 0.042 | 0.083 | -0.141 | -1.182 | 0.185 | -0.046 |
| -0.182 | -0.160 | -0.077 | 0.111 | 0.994 | 1.093 | 0.132 | 0.129 | 0.228 | -0.126 | -0.402 | 0.140 | 0.544 | -0.018 | 1.458 | 0.032 | -0.077 | 0.137 | 0.434 | -0.633 | 0.081 | -0.124 |
| 0.518 | -0.338 | -0.623 | -0.045 | 1.739 | 0.729 | 0.311 | -0.233 | 1.239 | -0.408 | -0.782 | 0.194 | -0.220 | 0.223 | 1.593 | -0.035 | -0.206 | 0.122 | 1.324 | 0.292 | 0.919 | -0.254 |
| 0.770 | -0.236 | -0.438 | -0.100 | 2.276 | 0.417 | 0.282 | -0.548 | 1.662 | -0.446 | -0.594 | 0.029 | -0.226 | -0.028 | 1.766 | -0.129 | -0.251 | 0.103 | 1.559 | 0.451 | 0.655 | -0.173 |
| 1.418 | -0.476 | -0.013 | -0.381 | 3.118 | 0.039 | 0.079 | -0.257 | 2.434 | -0.071 | -0.654 | -0.418 | -0.285 | -0.054 | 1.969 | -0.364 | -0.145 | -0.036 | 1.995 | 0.529 | 0.903 | 0.011 |
| 2.349 | 0.033 | 0.098 | 0.196 | 3.623 | -0.248 | 0.045 | 0.062 | 3.649 | -0.283 | 0.400 | 0.069 | -0.074 | -0.026 | 1.970 | -0.633 | -0.151 | -0.040 | 2.942 | 1.388 | -0.300 | -0.038 |
| 2.525 | 0.080 | 0.208 | 0.098 | 4.117 | -0.480 | 0.064 | 0.423 | 4.139 | -0.396 | 0.353 | -0.081 | 0.165 | -0.134 | 2.021 | -0.873 | -0.137 | 0.041 | 3.121 | 1.519 | -0.395 | 0.025 |
| -2.186 | 2.299 | 0.058 | 0.135 | -1.020 | -1.246 | 0.465 | 0.016 | -0.430 | -2.795 | 1.100 | -0.069 | 0.003 | 0.110 | -3.214 | 0.800 | -0.237 | -0.030 | 1.013 | 1.470 | -1.768 | -0.129 |
| -2.749 | -0.426 | 0.279 | -0.120 | -0.861 | -1.043 | 0.569 | 0.019 | -0.926 | -1.414 | -0.381 | -0.308 | 0.081 | 0.021 | -2.934 | 0.669 | -0.150 | -0.049 | 0.104 | -0.087 | -0.012 | -0.101 |
| -1.663 | -0.632 | 0.097 | 0.482 | -0.818 | -0.993 | 0.502 | 0.017 | -0.426 | -0.687 | -0.219 | 0.338 | -0.002 | -0.038 | -2.882 | 0.338 | -0.214 | 0.058 | 0.341 | -0.283 | -0.035 | -0.263 |
| -1.239 | -0.750 | 0.180 | 0.239 | -0.764 | -0.929 | 0.435 | 0.015 | -0.261 | -0.414 | -0.126 | 0.119 | 0.006 | -0.014 | -2.535 | 0.182 | -0.111 | 0.148 | 0.447 | -0.365 | -0.032 | -0.152 |
| -0.425 | -0.818 | -0.213 | 0.121 | -0.632 | -0.771 | 0.300 | 0.010 | -0.058 | -0.108 | -0.048 | 0.240 | -0.016 | -0.024 | -1.881 | -0.200 | 0.177 | -0.084 | 0.553 | -0.450 | -0.038 | -0.164 |
| 0.469 | -0.248 | -0.298 | 0.018 | -0.494 | -0.607 | 0.138 | 0.005 | 0.149 | 0.216 | 0.048 | 0.190 | -0.025 | -0.017 | -1.548 | -0.431 | 0.106 | -0.080 | 0.671 | -0.543 | -0.040 | -0.111 |
| 0.939 | -0.173 | 0.000 | -0.043 | -0.195 | -0.222 | 0.430 | 0.015 | 0.382 | 0.591 | 0.168 | 0.011 | -0.024 | 0.003 | -0.979 | -0.693 | 0.282 | 0.217 | 0.812 | -0.654 | -0.040 | -0.004 |
| 1.484 | 0.032 | 0.216 | 0.036 | -0.106 | -0.115 | 0.343 | 0.012 | 0.623 | 0.968 | 0.277 | -0.018 | -0.036 | 0.008 | -0.761 | -0.819 | 0.303 | 0.052 | 0.949 | -0.762 | -0.043 | 0.047 |
| 1.879 | 0.181 | 0.261 | -0.026 | 0.000 | 0.023 | 0.477 | 0.016 | 0.750 | 1.171 | 0.339 | -0.086 | -0.039 | 0.016 | -0.536 | -0.971 | 0.450 | 0.059 | 1.024 | -0.821 | -0.043 | 0.094 |
| 2.269 | 0.323 | 0.082 | -0.161 | 0.347 | 0.429 | -0.027 | -0.001 | 0.808 | 1.262 | 0.367 | -0.110 | -0.040 | 0.019 | -0.131 | -1.322 | -0.086 | -0.136 | 1.058 | -0.847 | -0.043 | 0.113 |
| 2.735 | 0.247 | -0.084 | 0.016 | 0.500 | 0.625 | 0.087 | 0.003 | 0.985 | 1.518 | 0.422 | 0.140 | -0.071 | -0.004 | 0.059 | -1.418 | -0.020 | -0.247 | 1.140 | -0.916 | -0.052 | 0.048 |
| 0.702 | 1.496 | -0.077 | 0.043 | -0.500 | -0.694 | -1.562 | -0.053 | | | | | | | 0.167 | -0.038 | -0.238 | -0.076 | -2.822 | 2.270 | 0.138 | 0.022 |
| | | | | -1.521 | -1.883 | 0.135 | 0.004 | | | | | | | | | | | -2.508 | 2.017 | 0.122 | 0.008 |
| | | | | -1.400 | -1.740 | -0.037 | -0.001 | | | | | | | | | | | -2.276 | 1.824 | 0.095 | -0.208 |
| | | | | -1.309 | -1.624 | 0.034 | 0.001 | | | | | | | | | | | -1.999 | 1.603 | 0.085 | -0.173 |
| | | | | -1.061 | -1.314 | 0.069 | 0.002 | | | | | | | | | | | -1.226 | 0.987 | 0.060 | 0.006 |
| | | | | -0.669 | -0.850 | -0.400 | -0.014 | | | | | | | | | | | -0.927 | 0.744 | 0.041 | -0.051 |
| | | | | -0.478 | -0.612 | -0.399 | -0.014 | | | | | | | | | | | -0.698 | 0.568 | 0.047 | 0.192 |
| | | | | -0.281 | -0.376 | -0.583 | -0.020 | | | | | | | | | | | -0.526 | 0.434 | 0.051 | 0.361 |
| | | | | -0.087 | -0.138 | -0.637 | -0.022 | | | | | | | | | | | -0.306 | 0.261 | 0.048 | 0.470 |
| | | | | 0.056 | 0.033 | -0.772 | -0.026 | | | | | | | | | | | -0.092 | 0.086 | 0.031 | 0.379 |
| | | | | 0.597 | 0.739 | -0.065 | -0.002 | | | | | | | | | | | 0.169 | -0.124 | 0.019 | 0.393 |
| | | | | 0.429 | 0.502 | -0.669 | -0.023 | | | | | | | | | | | | | | |

### PCA loading scores

| 45 days | | | | | | | | | | | | | | | | | | | | | |
|---|---|---|---|---|---|---|---|---|---|---|---|---|---|---|---|---|---|---|---|---|---|
| **Chamber Size** | | | | **Wall Geometry** | | | | **Cleaning Agent Scent** | | | | | | **Floor Texture** | | | | **Test Session Room** | | | |
| **PC1** | **PC2** | **PC3** | **PC4** | **PC1** | **PC2** | **PC3** | **PC4** | **PC1** | **PC2** | **PC3** | **PC4** | **PC5** | **PC6** | **PC1** | **PC2** | **PC3** | **PC4** | **PC1** | **PC2** | **PC3** | **PC4** |
| -3.376 | 1.625 | 0.113 | -0.281 | -2.060 | 3.334 | -0.460 | 0.313 | -5.037 | 0.901 | -0.811 | 0.500 | -0.179 | -0.004 | -1.062 | -0.101 | 0.262 | -0.194 | -3.965 | 1.604 | -0.228 | -0.501 |
| -2.735 | 1.390 | 0.631 | -0.427 | -1.516 | 2.758 | -0.151 | 0.711 | -3.925 | 0.166 | 0.465 | 0.518 | 0.591 | -0.048 | -0.528 | 0.030 | 0.590 | -0.061 | -3.348 | 1.718 | 0.124 | -0.175 |
| -1.041 | 0.322 | 0.655 | 0.513 | 0.278 | 2.078 | -0.411 | -0.099 | -2.297 | 0.433 | 0.860 | -0.732 | -0.153 | -0.019 | 0.714 | -0.328 | 0.103 | 0.023 | -1.549 | 1.397 | -0.249 | 1.139 |
| -0.015 | 0.064 | 0.129 | 0.097 | 0.758 | 1.530 | -0.376 | -0.044 | -0.205 | 0.454 | -0.233 | 0.001 | -0.355 | 0.276 | 1.078 | -0.258 | 0.046 | 0.016 | 0.504 | 0.517 | -0.026 | 0.006 |
| 0.741 | 0.112 | 0.361 | 0.303 | 1.893 | 0.768 | -0.170 | -0.551 | 0.515 | 0.843 | -0.043 | -0.124 | -0.257 | 0.373 | 1.861 | -0.365 | 0.041 | -0.102 | 1.196 | 0.760 | -0.039 | -0.053 |
| 1.000 | 0.154 | 0.465 | 0.402 | 2.135 | -0.088 | -0.151 | 0.210 | 1.294 | 0.861 | -0.194 | 0.112 | -0.055 | -0.321 | 2.154 | -0.025 | 0.029 | 0.006 | 1.394 | 0.892 | -0.059 | -0.058 |
| 1.284 | -0.206 | 0.502 | 0.274 | 2.379 | -0.295 | -0.018 | 0.079 | 1.811 | 0.673 | -0.087 | 0.171 | -0.278 | -0.333 | 2.320 | -0.034 | 0.105 | -0.057 | 1.891 | 0.603 | 0.110 | 0.166 |
| 1.624 | -0.231 | 0.366 | 0.101 | 2.576 | -0.514 | 0.039 | 0.071 | 2.558 | 0.482 | -0.208 | 0.410 | -0.057 | -0.152 | 2.467 | -0.011 | 0.122 | -0.077 | 2.556 | 0.360 | 0.213 | -0.281 |
| 2.376 | -0.015 | 0.089 | 0.220 | 3.432 | -1.804 | 0.049 | 0.291 | 3.676 | 0.552 | -0.488 | 0.437 | 0.469 | 0.267 | 3.099 | 0.229 | -0.100 | -0.151 | 3.634 | 0.272 | 0.068 | -1.136 |
| -2.468 | -1.843 | -0.438 | -0.115 | -2.380 | -2.710 | -0.599 | 0.087 | -0.157 | -3.029 | -0.169 | 0.246 | -0.168 | 0.013 | -2.874 | 2.019 | -0.323 | -0.244 | 0.067 | -1.786 | 0.244 | 0.696 |
| -1.873 | -2.355 | -0.175 | -0.184 | -2.117 | -3.320 | -0.367 | 0.861 | 0.712 | -3.631 | 0.749 | 0.046 | 0.008 | 0.008 | -2.162 | 2.207 | 0.396 | 0.114 | 0.720 | -1.962 | 0.479 | 1.321 |
| -2.081 | 0.507 | -0.142 | 0.710 | -1.427 | -0.541 | 0.306 | -0.583 | -0.594 | -0.950 | -0.810 | -0.032 | -0.009 | -0.019 | -3.106 | -1.807 | 0.542 | 0.060 | -0.198 | -0.361 | -0.342 | -0.076 |
| -0.811 | 0.143 | -0.754 | 0.036 | -0.613 | -0.230 | 0.142 | -0.248 | -0.410 | -0.678 | -0.632 | -0.116 | 0.017 | -0.018 | -2.041 | -1.784 | -0.134 | -0.170 | -0.023 | -0.124 | -0.341 | -0.013 |
| -0.195 | 0.223 | -0.792 | 0.444 | -0.457 | -0.167 | 0.130 | -0.181 | -0.094 | -0.284 | -0.563 | -0.563 | 0.141 | -0.035 | -1.523 | -0.656 | -0.209 | -0.173 | 0.270 | 0.193 | -0.640 | 0.088 |
| 0.708 | 0.577 | -0.700 | -0.208 | -0.430 | -0.157 | 0.120 | -0.171 | 0.136 | 0.138 | -0.076 | -0.328 | 0.088 | -0.015 | -0.902 | -0.271 | -0.518 | 0.178 | 0.495 | 0.589 | -0.304 | 0.173 |
| 0.978 | 0.600 | -0.810 | -0.204 | -0.153 | 0.002 | 0.324 | -0.010 | 0.219 | 0.243 | -0.054 | -0.441 | 0.119 | -0.019 | -0.683 | -0.209 | -0.550 | 0.224 | 0.572 | 0.673 | -0.378 | 0.200 |
| 1.321 | 0.568 | -0.929 | -0.324 | 0.094 | 0.086 | 0.228 | 0.083 | 0.296 | 0.352 | 0.008 | -0.491 | 0.134 | -0.020 | -0.268 | -0.307 | -0.542 | 0.179 | 0.644 | 0.767 | -0.392 | 0.226 |
| 2.207 | 0.722 | -0.577 | -0.307 | 0.375 | 0.147 | -0.054 | 0.158 | 0.697 | 0.964 | 0.457 | -0.596 | 0.172 | -0.014 | 0.367 | -0.087 | -0.528 | 0.016 | 1.026 | 1.306 | -0.313 | 0.364 |
| 0.654 | -0.115 | 0.851 | -0.314 | -0.679 | 0.038 | 1.581 | -0.016 | 0.310 | 0.582 | 0.704 | 0.378 | -0.087 | 0.031 | 0.149 | 0.146 | 0.128 | 0.070 | -0.539 | -0.064 | 2.465 | -0.157 |
| 1.065 | -0.251 | 1.349 | -0.508 | -0.308 | 0.266 | 1.929 | 0.213 | 0.494 | 0.927 | 1.122 | 0.603 | -0.139 | 0.049 | 0.517 | 0.092 | 0.611 | 0.304 | -0.339 | 0.281 | 2.737 | -0.081 |
| 0.236 | -0.740 | -0.072 | -0.085 | -1.731 | -0.705 | 0.133 | -0.750 | | | | | | | 0.137 | 0.490 | -0.023 | 0.013 | -1.782 | -2.406 | 0.040 | -0.640 |
| 0.399 | -1.250 | -0.122 | -0.143 | -0.671 | -0.297 | -0.064 | -0.312 | | | | | | | 0.288 | 1.030 | -0.049 | 0.027 | -0.943 | -1.294 | -0.059 | -0.340 |
| | | | | -0.561 | -0.259 | -0.107 | -0.270 | | | | | | | | | | | -0.857 | -1.187 | -0.096 | -0.309 |
| | | | | -0.145 | -0.182 | -0.587 | -0.171 | | | | | | | | | | | -0.543 | -0.902 | -0.610 | -0.204 |
| | | | | -0.008 | -0.139 | -0.661 | -0.124 | | | | | | | | | | | -0.437 | -0.777 | -0.680 | -0.167 |
| | | | | 0.220 | -0.058 | -0.735 | -0.035 | | | | | | | | | | | -0.258 | -0.550 | -0.739 | -0.104 |
| | | | | 0.411 | -0.001 | -0.851 | 0.029 | | | | | | | | | | | -0.110 | -0.379 | -0.852 | -0.052 |
| | | | | -0.065 | -0.042 | -0.072 | -0.042 | | | | | | | | | | | -0.223 | -0.405 | -0.378 | -0.086 |
| | | | | 0.056 | 0.036 | 0.062 | 0.036 | | | | | | | | | | | -0.121 | -0.220 | -0.206 | -0.047 |
| | | | | 0.214 | 0.139 | 0.236 | 0.138 | | | | | | | | | | | 0.012 | 0.021 | 0.020 | 0.004 |
| | | | | 0.502 | 0.326 | 0.554 | 0.325 | | | | | | | | | | | 0.254 | 0.462 | 0.431 | 0.098 |

### Explained Variance

| 2 days | | | | | | | | | | | | | | | | | | | | | |
|---|---|---|---|---|---|---|---|---|---|---|---|---|---|---|---|---|---|---|---|---|---|
| **Chamber Size** | | | | **Wall Geometry** | | | | **Cleaning Agent Scent** | | | | | | **Floor Texture** | | | | **Test Session Room** | | | |
| **PC1** | **PC2** | **PC3** | **PC4** | **PC1** | **PC2** | **PC3** | **PC4** | **PC1** | **PC2** | **PC3** | **PC4** | **PC5** | **PC6** | **PC1** | **PC2** | **PC3** | **PC4** | **PC1** | **PC2** | **PC3** | **PC4** |
| 1.575 | 0.972 | 0.699 | 0.292 | 1.420 | 1.187 | 0.686 | 0.324 | 1.748 | 1.384 | 0.881 | 0.409 | 0.233 | 0.182 | 1.853 | 0.710 | 0.214 | 0.131 | 1.406 | 1.144 | 0.737 | 0.413 |
| 0.620 | 0.236 | 0.122 | 0.021 | 0.504 | 0.352 | 0.118 | 0.026 | 0.509 | 0.319 | 0.129 | 0.028 | 0.009 | 0.006 | 0.858 | 0.126 | 0.011 | 0.004 | 0.494 | 0.327 | 0.136 | 0.043 |
| 0.620 | 0.856 | 0.979 | 1.000 | 0.504 | 0.856 | 0.974 | 1.000 | 0.509 | 0.828 | 0.957 | 0.985 | 0.994 | 1.000 | 0.858 | 0.984 | 0.996 | 1.000 | 0.494 | 0.821 | 0.957 | 1.000 |

| 28 days | | | | | | | | | | | | | | | | | | | | | |
|---|---|---|---|---|---|---|---|---|---|---|---|---|---|---|---|---|---|---|---|---|---|
| **Chamber Size** | | | | **Wall Geometry** | | | | **Cleaning Agent Scent** | | | | | | **Floor Texture** | | | | **Test Session Room** | | | |
| **PC1** | **PC2** | **PC3** | **PC4** | **PC1** | **PC2** | **PC3** | **PC4** | **PC1** | **PC2** | **PC3** | **PC4** | **PC5** | **PC6** | **PC1** | **PC2** | **PC3** | **PC4** | **PC1** | **PC2** | **PC3** | **PC4** |
| 1.847 | 0.687 | 0.272 | 0.205 | 1.513 | 1.219 | 0.445 | 0.170 | 2.194 | 0.947 | 0.439 | 0.218 | 0.201 | 0.102 | 1.778 | 0.886 | 0.204 | 0.113 | 1.495 | 1.243 | 0.427 | 0.189 |
| 0.853 | 0.118 | 0.019 | 0.011 | 0.572 | 0.371 | 0.049 | 0.007 | 0.802 | 0.149 | 0.032 | 0.008 | 0.007 | 0.002 | 0.790 | 0.196 | 0.010 | 0.003 | 0.559 | 0.387 | 0.046 | 0.009 |
| 0.853 | 0.971 | 0.990 | 1.000 | 0.572 | 0.943 | 0.993 | 1.000 | 0.802 | 0.952 | 0.984 | 0.992 | 0.998 | 1.000 | 0.790 | 0.986 | 0.997 | 1.000 | 0.559 | 0.946 | 0.991 | 1.000 |

| 45 days | | | | | | | | | | | | | | | | | | | | | |
|---|---|---|---|---|---|---|---|---|---|---|---|---|---|---|---|---|---|---|---|---|---|
| **Chamber Size** | | | | **Wall Geometry** | | | | **Cleaning Agent Scent** | | | | | | **Floor Texture** | | | | **Test Session Room** | | | |
| **PC1** | **PC2** | **PC3** | **PC4** | **PC1** | **PC2** | **PC3** | **PC4** | **PC1** | **PC2** | **PC3** | **PC4** | **PC5** | **PC6** | **PC1** | **PC2** | **PC3** | **PC4** | **PC1** | **PC2** | **PC3** | **PC4** |
| 1.632 | 0.921 | 0.615 | 0.334 | 1.383 | 1.282 | 0.579 | 0.329 | 1.966 | 1.253 | 0.559 | 0.415 | 0.235 | 0.168 | 1.735 | 0.919 | 0.353 | 0.144 | 1.470 | 1.021 | 0.768 | 0.456 |
| 0.666 | 0.212 | 0.095 | 0.028 | 0.478 | 0.411 | 0.084 | 0.027 | 0.644 | 0.261 | 0.052 | 0.029 | 0.009 | 0.005 | 0.752 | 0.211 | 0.031 | 0.005 | 0.540 | 0.260 | 0.147 | 0.052 |
| 0.666 | 0.878 | 0.972 | 1.000 | 0.478 | 0.889 | 0.973 | 1.000 | 0.644 | 0.905 | 0.957 | 0.986 | 0.995 | 1.000 | 0.752 | 0.964 | 0.995 | 1.000 | 0.540 | 0.801 | 0.948 | 1.000 |

### Eigenvalues

| 2 days | | | | | |
|---|---|---|---|---|---|
| **PC** | **Chamber Size** | **Wall Geometry** | **Cleaning Agent Scent** | **Floor Texture** | **Test Session Room** |
| PC1 | 2.481 | 2.016 | 3.055 | 3.433 | 1.977 |
| PC2 | 0.944 | 1.409 | 1.914 | 0.504 | 1.309 |
| PC3 | 0.489 | 0.470 | 0.775 | 0.046 | 0.544 |
| PC4 | 0.085 | 0.105 | 0.168 | 0.017 | 0.170 |
| PC5 | | | 0.054 | | |
| PC6 | | | 0.033 | | |
| 28 days | | | | | |
| **PC** | **Chamber Size** | **Wall Geometry** | **Cleaning Agent Scent** | **Floor Texture** | **Test Session Room** |
| PC1 | 3.412 | 2.288 | 4.813 | 3.161 | 2.236 |
| PC2 | 0.472 | 1.485 | 0.896 | 0.784 | 1.546 |
| PC3 | 0.074 | 0.198 | 0.193 | 0.042 | 0.182 |
| PC4 | 0.042 | 0.029 | 0.048 | 0.013 | 0.036 |
| PC5 | | | 0.040 | | |
| PC6 | | | 0.010 | | |
| 45 days | | | | | |
| **PC** | **Chamber Size** | **Wall Geometry** | **Cleaning Agent Scent** | **Floor Texture** | **Test Session Room** |
| PC1 | 2.663 | 1.913 | 3.864 | 3.009 | 2.162 |
| PC2 | 0.848 | 1.643 | 1.569 | 0.845 | 1.041 |
| PC3 | 0.378 | 0.336 | 0.312 | 0.125 | 0.589 |
| PC4 | 0.111 | 0.109 | 0.172 | 0.021 | 0.208 |
| PC5 | | | 0.055 | | |
| PC6 | | | 0.028 | | |

**Supplemental Table 3.** Tables **describing** PCA loading scores, Explained Variances and Eigenvalues for each interval (parameter variables vs. sex). "QA": quaternary ammonium.

#### Interpreting the multivariate analysis

- As shown in the **Heatmaps** in **Figure S-3 A**, strong positive correlations dominate when comparing the four selected contexts, providing no useful information to interpret the relative weights of the parameters. In **Figure S3-B**, however, varying all 11 variants of the 5 parameters at once, show a few differences between females and males at 2 and 28 days, but a noticeable change at 45 days in the correlation matrices, especially in males, compared to females. **Figure S-3 C shows** a few negative correlations on the reactions between grid vs. smooth FLOOR textures at 28 days, and WALL geometry at 45 days.

- **Figure 4-S A** shows the PCA (Loadings) plot for the four contexts, organized by sex (columns) and time (rows). PC2 appears to be more representative of these variations, while projections along PC1 are only on the positive side. At 2 days, contexts A, B, and C are clustered and orthogonal to D for both sexes, indicating a positive correlation among them and independence from the response to D. At 28 days, context C is independent, uncorrelated with the other three, for both sexes. A distinct pattern emerges only at 45 days, when females show a high correlation between contexts CD and AB, while males show high correlation only between B and D.

- In **Figure 4-S B**, 11 variables corresponding to five behavioral parameters were analyzed in blocks, revealing several sex-specific patterns: in females, at 2 days, four variables associated with the conditioning/training context A (excluding SCENT) clustered together (positively correlated) in the upper left quadrant and were orthogonal, i.e., statistically independent, from the novel context variables, which is to be expected. The exception was the non-training context variable "smooth" – wherein males do not discriminate between grid and smooth

FLOORS at 2 days and females failed to discriminate at any timepoint. For females, the clustering of context A variables dispersed at 28 days and partially regrouped at 45 days, now in the upper right quadrant. In males, a similar clustering was observed at 2 days, although with evident confounding between FLOOR and SIZE variables. Unlike females, however, variables are dispersed for males at 28 days – with a negative correlation between grid and smooth FLOOR, and recluster at 45 days with the context A variables now predominantly distributed within the negative PC2 quadrants. Initially, in these plots the three distinct SCENTs are clustered together, positively correlated, however in more remote memory ethanol became orthogonal to the other odors, an independency that suggests that both sexes continue to associate this odor with the original aversive experience up to 45 days later.

- In **Figure S-4 C**, we analyzed the variables separated by parameter to identify sex-specific and time-dependent patterns:
    - In relation to FLOOR, sexes responded independently (orthogonally) at 2 days, with neither group clearly distinguishing between textures. At later timepoints, males tend to cluster with females, although consistently maintaining independent behavior for one variable - grid FLOOR at 28 days, and smooth FLOOR at 45 days. Females, on the other hand, do not distinguish FLOOR textures well at any timepoint.;
    - both sexes respond similarly to planar or curved WALLs up to 28 days, although these two variables are orthogonal, i.e., vary independently. This pattern is disrupted at 45 days, when males tend to distinguish planar from curved WALLs (vectors forming obtuse angles), whereas females tend to cluster these variables, responding similarly to both geometries;
    - In relation to chamber SIZE, subjects responded to the larger ones independently at all times. Specifically, females exhibited independent responses at 2 and 28 days, and males at 45 days.
    - similarly to what was observed with WALLs, subjects responded the same way for each ROOM, independent of sex, with the conditioning

ROOM and novel ROOM orthogonal to each other at 2 and 28 days (the only difference being the quadrant of these vectors). At 45 days, however, this pattern is disrupted, with males distinguishing ROOMs well (vectors forming obtuse angles), whereas females tend to cluster these variables, possibly confounding them;

- The most notable aspect in the analysis of these three distinct SCENTs is that males and females clustered similarly to ethanol – odor associated with the training context - while maintaining independency (orthogonality) from the other typically clustered odors, at least at 2 and 28 days. At 45 days, there is a subtle tendency of females to respond distinctly to quaternary ammonium, and the influence of ethanol tends to diminish at 45 days.
